# Phylogenomics reveals incomplete lineage sorting and ancient hybrid drove the radiation of macaques

**DOI:** 10.1101/2023.01.09.523240

**Authors:** Xinxin Tan, Jiwei Qi, Zhijin Liu, Pengfei Fan, Gaoming Liu, Liye Zhang, Ying Shen, Jing Li, Christian Roos, Xuming Zhou, Ming Li

**Affiliations:** CAS Key Laboratory of Animal Ecology and Conservation Biology, Institute of Zoology, Chinese Academy of Sciences, Beijing 100101, China; Geneplus-Beijing Institute, Beijing 102206, China; College of Life Sciences, Capital Normal University, Beijing 100049, China; School of Life Sciences, Sun Yat-Sen University, Guangzhou 510275, China; Primate Genetics Laboratory, German Primate Center, Leibniz Institute for Primate Research, Kellnerweg 4, 37077, Göttingen, Germany; Key Laboratory of Bio-resources and Eco-environment of Ministry of Education, College of Life Sciences, Sichuan University, Chengdu 610064, China; Gene Bank of Primates, German Primate Center, Leibniz Institute for Primate Research, Kellnerweg 4, 37077, Göttingen, Germany

**Keywords:** *Macaca*, phylogenomics, incomplete lineage sorting, hybridization.

## Abstract

The genus *Macaca*, with 23 species assigned into four to seven species groups, exhibits the largest geographic range and represents the most successful adaptive radiation of nonhuman primates. Here, we conducted phylogenomic analyses of 16 newly generated and eight published macaque genomes and found a strong support for the division of this genus into seven species groups. Both ancient hybrid and incomplete lineage sorting (ILS) have contributed to the radiation and evolution of macaques. Particularly, the contradicting phylogenetic positions among *silenus/nigra, fascicularis/mulatta* and *arctoides*/*sinica* lineages were likely resulted from high level of ILS and potential hybridization between the ancestors of the *arctoides*/*sinica* and *silenus/nigra* lineages. Furthermore, an integrated scenario for macaque radiation is reconstructed by the help of the dated phylogenetic tree combined with documented history records. This study provides insights into ancient introgression involved in the radiation of macaques, which may help us to understand the rapid speciation of nonhuman primates.

## Introduction

Introgression and hybridization are important driving forces in speciation and evolution, and over the past decade increasing attention has been paid to their role in evolutionary diversification [1, 2, 3, 4, 5, 6, 7]. Hybridization has been recognized as one of the most interesting topics in evolutionary biology, and the genetic introgression, caused by hybridization, is not always maladaptive and could be a potent evolutionary force [3, 8, 9, 10]. With the development of high-throughput sequencing technology and new computational tools, phylogenomics emerged and allows researchers to explicitly account for hybridization when constructing species trees [11, 12]. However, incomplete lineage sorting (ILS), an alternative explanation for contradicting gene trees, needs to be considered as well when reconstructing a species tree [13]. ILS occurs mainly during rapid speciation events, when new lineages descend within a short time period from ancestors, particularly when these have large effective population sizes (*Ne*) [14]. As a result, ILS will cause ancestral polymorphisms to persist even when descendant lineages diverged [15]. Therefore, both ILS and gene flow or hybridization can lead to topological discordance between gene trees and species trees [16, 17]. For example, Edelman et al. [18] used 20 *de novo* genome assemblies to explore the speciation history and gene flow in rapidly radiating *Heliconius* butterflies and distinguish ILS from introgression. They discovered that introgressed loci are underrepresented in low-recombination and gene-rich regions [18]. For primates, information about a genome-wide view on hybridization is still limited but genome sequencing efforts are on the way [19]. For example, Rogers, et al. [20] confirmed reticulation in baboons and documented multiple episodes of admixture and introgression during the radiation of *Papio* using a comparative genomic approach.

The genus *Macaca* (Primates: Cercopithecidae) exhibits the largest geographic range and represents the most successful adaptive radiation of nonhuman primates [21] (**Fig. 1A**). The 23 currently recognized macaque species are subdivided into four to seven species groups according to similarities in morphology, ecology, behavior, distribution and genetics [21, 22, 23, 24, 25, 26, 27, 28, 29, 30, 31, 32, 33] (**supplementary Fig. S1**). However, the number and composition of the species groups is disputed. Before the whole-genome age, phylogenetic analyses were mainly based on a few mitochondrial and/or nuclear markers resulting in contradicting phylogenetic relationships [26, 34, 35, 36], which are partially caused by introgression and hybridization events among various macaque species.

**Figure 1.**
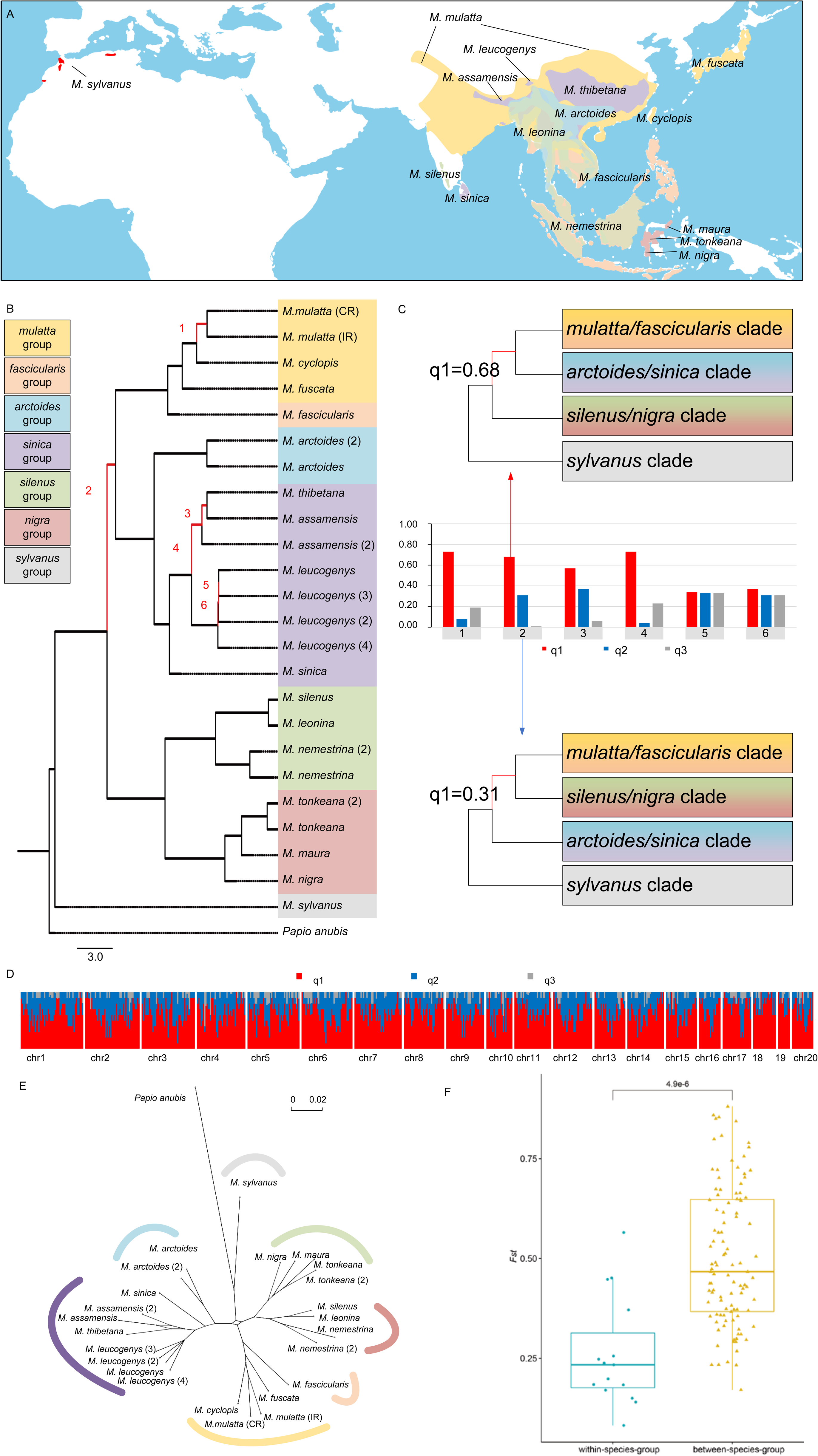
Macaque phylogenetics and differentiation. (A) Distribution ranges of the 16 macaque species investigated in this study. The range overlaps are shown in blended colors (orange, purple, green, blue and yellow). (B) A MSC species tree was constructed from 6,351 individual genome fragments. Branches with low q1 (<0.8) are numbered 1 to 6. All branches receive maximal support (P = 1.0, ASTRAL analysis). Branch lengths were calculated from a ML analysis. Different species groups are represented by different background colors which are show to the left. (C) ASTRAL quartet-score analyses for branches 1 to 6 in Fig. 1B. Quartet scores were calculated for the three possible arrangements (q1 to q3) for the respective branch. The principal quartet trees are depicted, with q1 representing the species tree. Two alternative topologies for branch no. 2 as shown in Fig. 1C. (D) Genealogical discordance across the genome of rhesus macaque, demonstrated by a window analysis (200-kb window size) of a full-genome alignment mapped to Panu_3.0 (chromosomes showed at the bottom). The y axis indicates the percentage of windows within a given interval (every 5 windows next to each other) that conform to (red) or reject (blue and grey) the species tree. (E) Consensus networks for macaques from 6,351 genome fragments with a minimum threshold of 30% to form an edge. (F) Distributions of *F_ST_* within (blue) and between (yellow) different macaque species groups. The exact *P* value of above were computed based on Wilcoxon test.

Introgression and hybridization between macaque species are common, for instance, among Sulawesi macaques, between *M. mulatta* and *M. fascicularis*, and between *M. mulatta* and *M. thibetana* [37, 38, 39, 40, 41, 42, 43, 44, 45]. Moreover, for *M. arctoides* a hybrid origin was suggested as different molecular markers place the species into different clades [33, 34, 35, 36, 46]. However, most analyses have focused on gene flow within macaque species groups, or between species of the *fascicularis*/*mulatta* and *arctoides*/*sinica* species groups. Few studies included samples from all species groups when identifying genus-wide hybridization, and few attempted to detect ancient hybridization events in macaques [36, 47]. A recent study reported the history of the genus *Macaca* using 14 individuals from nine species and detected extensive gene flow signals, with the strongest signals between the *fascicularis/mulatta* and *silenus* groups [48]. Introgression signals among island species, such as Sulawesi macaques and other species, were also observed. This put forward a variety of possible reasons for this phenomenon including the genomic similarity of closely related species or ancestral introgression. Therefore, whether hybridization occurred among all macaques, especially among species that are geographically distant still needs to be investigated. Furthermore, it remains unclear how much ancient hybridization or ILS have contributed to the diversity of the macaque genome.

In this study, we aimed to trace the complex, likely reticulated evolutionary history of macaques and newly sequenced whole genomes of 16 individuals representing 13 Asian (*M. mulatta, M. cyclopis, M. fuscata, M. arctoides, M. assamensis, M. leucogenys, M. sinica, M. leonina, M. silenus, M. nemestrina, M. maura, M. tonkeana, M. nigra*) and the only African macaque species (*M. sylvanus*). The sequences were mapped to the latest *Papio anubis* reference genome (Panu_3.0) to obtain single nucleotide variants (SNVs). Combined with eight published macaque genomes, we performed a comprehensive genomics analysis among 24 macaque individuals, representing 16 species and all species groups, aiming to reconstruct the phylogeny of the genus *Macaca,* investigate ILS and hybridization among macaque species, and trace the evolutionary history and rapid radiation of the genus *Macaca*.

## Results

### Genome Sequencing and Variant Discovery

Whole genome sequencing was performed with the Illumina HiSeq 2500 and 4000 platforms. In total, more than 1,364 Gbp clean data (quality-controlled reads) were obtained and the statistics results of the clean data for each species are shown in **supplementary table S1**. Published genome data of eight macaques (*M. mulatta* (Chinese rhesus macaque, CR), *M. mulatta* (Indian rhesus macaque, IR), *M. fascicularis*, *M. arctoides*, *M. assamensis, M. thibetana, M. nemestrina* and *M. tonkeana*) were downloaded from NCBI (**supplementary table S1**) and two of them overlapped with [48]. Clean data for each species were mapped to the *Papio anubis* reference genome (Panu_3.0) and yielded genome-wide coverage from 16.85 to 42.34× with an average effective depth at 28.24× (**supplementary table S1**). We picked a more distant reference genome (*Papio anubis*) rather than the more recently evolved macaque (*Macaca mulatta*) to avoid biases that would arise from mapping to an in-group reference genome. Even though this decision likely led to some degree of lost data (in regions that lacked ortholog to *Papio anubis*), however, the mapping coverage (4×) for macaques range from 93.55% to 97.59%, which is very similar to mapping coverage (4×) for *Papio* (98.75%), suggesting a potential bias to be limited. The mapping rate for *M. nemestrina* (2) is lower than 90%, which could be caused by low DNA quality.

The number of high-quality SNVs for each individual ranged from 15,943,244 to 21,778,812, counting both homozygous and heterozygous alternatives (**supplementary table S2**). Genome-wide heterozygosity varied considerably among macaques (**supplementary Fig. S2** and **supplementary table S2**) which may partially reflect different demographic histories. Based on gene annotations from the reference genome, the exonic SNVs in macaques ranged from 225,336 to 257,712 (**supplementary tables S3** and **S4**). The species-specific SNVs for each macaque ranged from 1,279,649 to 8,688,344 (**supplementary table S2**).

### Phylogeny and divergence among macaques

In order to reconstruct the phylogeny of macaques, we generated consensus sequences of all genomes for each macaque individual. We split the aligned chromosomes into 6,351 non-overlapping genome fragments (GFs), each 200-kilo basepair (kbp) long, which representing 46% of the genome sequence. The multispecies coalescent (MSC) species tree based on the 6,351 GFs was supported with posterior probabilities (PPs) of 1.0 for all branches (**Fig. 1B** and **supplementary Fig. S3**). This species tree, consistent with neighbor joining (NJ) tree and maximum-likelihood (ML) tree based on autosomal SNVs (**supplementary Fig. S4** and **supplementary Fig. S5**), supported the segregation of macaques into seven well-supported clades or lineages that correspond to the classification of the genus into seven species groups [21, 29]. Among macaques, the African *sylvanus* lineage diverged first, followed by an initial separation of extant Asian macaques into a clade with the *silenus* and *nigra* lineages and a clade including all other Asian lineages. The latter segregated into a clade including the *arctoides* and *sinica* lineages and another clade with the *fascicularis* and *mulatta* lineages (**Fig. 1B** and **supplementary Figs. S3-S5**). The ML tree based on mitochondrial genomes (mitogenomes), consistent with previous results [32, 33], showed that among Asian macaques, the *silenus/nigra* clade separated first, followed by the *sinica* lineage, while among the remaining lineages, *fascicularis* split first and *mulatta* and *arctoides* afterwards (**supplementary Fig. S6** and **supplementary table S5**).

The quartet score ‘q’, which is the support value for possible phylogenetic arrangements, identified a conflict in resolving the branch leading to the ancestor of the *fascicularis/mulatta* and *arctoides/sinica* groups (**Fig. 1C-D**, branch no. 2 in **Fig. 1B** and **supplementary Fig. S3**). In addition, the topology cluster found that these two alternative topologies are most common among all 6,351 GF trees and represented by the colors red and blue, respectively in **Fig. 3A-B**. However, the reason for this conflict is unclear.

A consensus network analysis [40] of the GF trees recovered a network with bifurcating branches that emerged from a central cycle in the center of the network that indicates conflicting signals for the position of the *silenus/nigra* clade among Asian macaques (**Fig. 1E** and **supplementary Fig. S7**). At a threshold for conflicting edges of 30%, the position of the *silenus/nigra*, *arctoides/sinica* and *fascicularis/mulatta* clades were unascertainable (**supplementary Fig. S7**). When the threshold for conflicting edges was reduced, the phylogenetic signal became even more complex, indicating additional phylogenetic conflict and extensive gene flow in the data set (**supplementary Fig. S7**). *F_ST_* analyses suggested significant differences within (medians for *F_ST_* are 0.199) and between (medians for *F_ST_* are 0.467) different macaque groups (*P* < 0.0001) (**supplementary table S6** and **Fig. 1F**). A clear division of macaques into different clusters was also obtained by a principal component analysis (PCA) (**supplementary Fig. S8** and **supplementary table S7**).

The genus *Macaca* was estimated to have originated ca. 7.76 million years ago (Mya) during the Late Miocene using MCMCTREE in PAML v4.9 [49] (**Fig. 2** and **supplementary Fig. S9**). We revealed a divergence time for the split of *M. sylvanus* from the Asian macaques at 5.29 Mya (95% highest posterior density, HPD, 3.65–6.70), close to the Messinian Salinity Crisis [50]. In Asia, divergence of species groups occurred in the Pliocene, starting with the split between the *silenus/nigra* clade and the other four species groups 4.01 Mya (95% HPD 5.59–2.61), followed shortly afterwards by the separation of the ancestors of the *fascicularis/mulatta* and *arctoides/sinica* clades (3.79 Mya, 95% HPD 5.28–2.50). These three clades finally diverged into species groups 3.20 Mya (95% HPD 4.79–2.07; *silenus* and *nigra* groups), 3.01 Mya (95% HPD 4.52–1.84; *sinica* and *arctoides* groups) and 2.94 Mya (95% HPD 4.25–1.52; *mulatta* and *fascicularis* groups). Speciation within species groups took place in the Pleistocene, 2.56–1.39 Mya.

**Figure 2.**
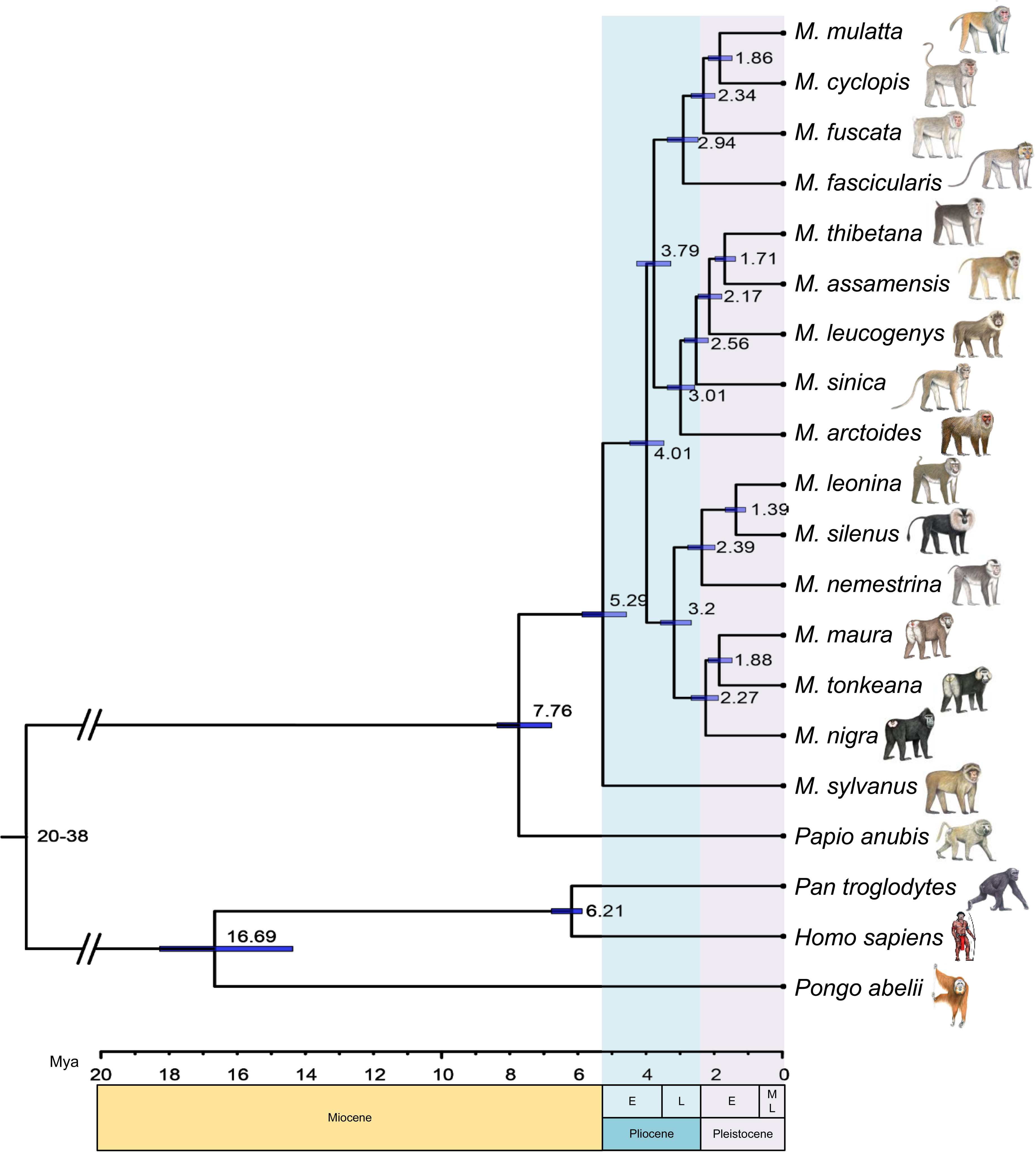
Divergence time tree of genus *Macaca*, estimated from 3,319,411 4d sites from 10,641 single-copy genes. Blue bars indicate 95% HPDs of divergence times and the time scale below shows million years ago (Mya). Abbreviations: L = late, M = middle, E = early. Four fossil-based calibration points are as follows: Catarrhini 20–38 Mya, Hominidae 13–18 Mya, *Homo*-*Pan* 6–7 Mya and *Papio*-*Macaca* 5–8 Mya.

### Contributions of ILS to Gene Tree Heterogeneity

Our study was aimed at investigating the extent to which levels of gene tree heterogeneity found in macaques can be attributed to incomplete lineage sorting (ILS), especially for the conflict among the *fascicularis/mulatta*, *arctoides/sinica* and *silenus/nigra* clades showed in **Fig. 1C** and **Fig. 3A-B**. To differentiate between ILS and introgression, which are both potential causes of phylogenetic discordance, we utilized QuIBL, a recently developed tree-based method introduced by Edelman et al. [18]. QuIBL estimates internal branch length distribution in discordant topologies for triplets of species and calculates the likelihood that this distribution is consistent with either ILS and introgression or ILS alone. Since this method is sensitive to recombination [18], we generated 3,175 trees based on small, non-overlapping windows (20 kb) separated by 400-kb steps. Our study revealed that within the *fascicularis/mulatta*, *silenus/nigra*, and *sinica/arctoides* clades, the Bayesian information criterion (BIC) test determined that 60.8% (73 out of 120 triplets) of triplets with internal branches of species exhibited phylogenetic discordances solely due to incomplete lineage sorting (ILS) (ΔBIC>10). Furthermore, only 9.1% (11 out of 120 triplets) of our tested triplets showed strong evidence of introgression (ΔBIC<-10) (**supplementary table S8**). For instance, using QuIBL on the triplet *M. leonina_M. mulatta_M. thibetana*, we inferred that only 1.33% of loci across the entire genome are introgressed. Moreover, we discovered that a mere 0.29% of genetic loci supported discordant topologies and were introgressed, indicating that interspecific introgression was limited among the species examined (**Fig. 3F, supplementary table S8).** Consequently, QuIBL analysis suggests that incomplete lineage sorting, rather than post-speciation introgression, is the predominant factor underlying the phylogenetic discordance among these species.

**Figure 3.**
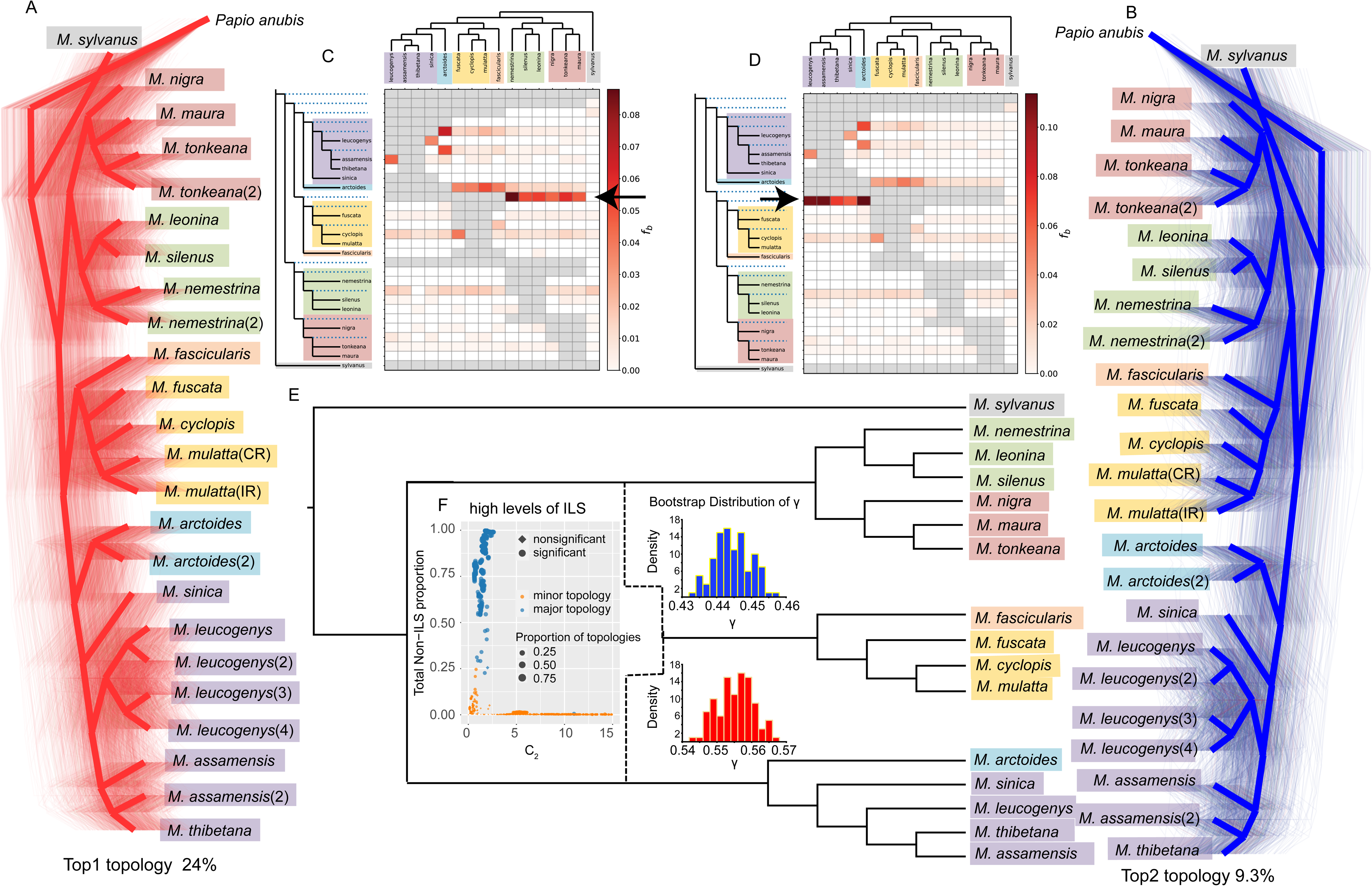
High levels of ILS and footprints of introgression contribute to gene tree heterogeneity. (A), (B) All samples’ DensiTree plots were created using 200 bp window trees spanning the whole genome. The top two most common topologies are represented by the colors red (A) and blue (B), respectively. (C), (D) The top two most popular topologies are taken as the reference trees, and a heatmap is shown to illustrate the statistical support for introgression between species pairs inferred from the Dsuit software. The Fbranch function was used to further process the Dtrios results for every branch on the tree, The black arrows refer to the infiltration of the *fascicularis/mulatta* clade with the *silenus/nigra* (C) and *sinica/arctoides* (D) clades, respectively. (E) The diagram illustrates the complex evolutionary relationships between macaque species, highlighting instances of hybridization and incomplete lineage sorting (ILS). The *fascicularis/mulatta* clade have a hybrid origin, while the reticulated patterns on the blue (*silenus/nigra*) and red (*sinica/arctoides*) γ correspond to the estimated probabilities of inheritance from the respective ancestral clades. (F) QuIBL results for the triplets. Relationships between internal branch length in coalescent units (C_2_) and total proportion of introgressed loci, blue symbols represent triplets with true topology, orange symbols represent triplets with discordant topology. Total proportion of introgressed loci was obtained by multiplying the probability that each topology corresponds to introgression by its genomic frequency.

### Contributions of hybrid to Gene Tree Heterogeneity

Previous studies suggested a vital role of hybridization in shaping the evolutionary history of macaques [35, 36, 37, 38, 41, 42, 43, 44, 45, 46, 51]. In this study, we set out to infer footprints of ancient introgression across macaques. To this end, we first applied the approach based on D-statistics [52] implemented in *Dsuite* [53] to detect introgression and hybridization (**Fig. 3A-B, supplementary table S9)**. This approach estimates D and f4 statistics for all possible combinations of trios in macaques based on two alternative topologies described in **Fig. 3A-B**, respectively, and then performs an f-branch test to assign gene flow to specific internal branches (**Fig. 3C-D)**. The f-branch test based on the species tree suggested an event of introgression between the ancestor of *fascicularis/mulatta* and the ancestor of *silenus/nigra* lineages **(Fig. 3C)**. While the f-branch test based on another topology suggested an event of introgression between the ancestor of *fascicularis/mulatta* and the ancestor of *arctoides/sinica* lineages **(Fig. 3D)**. The above results suggest that the genetic background of *fascicularis/mulatta* lineage is highly heterogeneous and the ancestor of *fascicularis/mulatta* lineage may be a hybrid lineage. In addition, ancient hybrid speciation was also be studied by topology-based ML analysis using PhyloNet [54], which identifies a reticulate node in the scenarios among the ancestor of *fascicularis/mulatta, arctoides/sinica* and *silenus/nigra* lineages (**supplementary Fig. S10**).

Next, we corroborated the evidence for hybridization of the ancestor of *fascicularis/mulatta* in HyDe [55], which can examine genome-scale data for a large number of taxa and detect the population that may have arisen through hybrid speciation, as well as its putative parental populations, by estimates the inheritance parameter (γ value)_ to quantify the genomic contributions of the parents to the hybrid. In this study, HyDe was used to verify for significant *p-*values in all combinations of the four taxa, which included one outgroup (*P. anubis*) and a triplet of in-group taxa. However only one of the four significant hybridization events showed a γ value as 0.55 (Z–score =306.84; *P* = 0), which indicated that the *fascicularis/mulatta* lineage is a hybrid speciation and the genomic contributions of its parents (*arctoides/sinica* and *silenus/nigra* lineages) is about 55% vs 45% (**supplementary table S11**). We further verified the hybridization for *fascicularis/mulatta* lineage at the individual level and found that all 5 samples showed a γ value as ranges from 0.534 to 0.571, which further declared a similar, but more nuanced, characterization of high levels of hybridization for *fascicularis/mulatta* lineage (**supplementary table S11)**. Finally, bootstrap resampling of individuals within hybrid populations were performed to obtain a distribution of gamma values to assess heterogeneity in levels of introgression (**Fig. 3F**).

Finally, *F*_d_ statistic [56] was initially used to calculate the potential introgression in 10-kb windows firstly and genome regions with top 0.05 high *F*_d_ values were selected. Given it is of challenge to accurately identify the introgressed regions in macaque genomes, we applied a two-layer hidden Markov model [57] to infer local ancestry of admixed individuals of the *fascicularis/mulatta* lineage. We selected the top 1% windows with high sum of the proportion of source population (*silenus/nigra* lineage) ancestry for all SNPs in a 10 kb windows with a high *F*_d_ values as the final potential introgression regions (**Supplementary tables S12**) and found some functional genes (e.g., *CERS3*, *ACER3*, *GSK3B*) present in these potential introgressed regions (**supplementary Fig. S11** and **supplementary table S13**). We then reconstructed a phylogenetic tree based on autosomal SNVs including only SNVs located in all putatively introgressed regions for *fascicularis/mulatta* lineage form *silenus/nigra* lineage and there is no doubt that the *arctoides/sinica* lineages diverged first among Asian macaques (**supplementary Fig. S12**). In addition, we reconstructed a phylogenetic tree excluding SNVs located in all putatively introgressed regions for *fascicularis/mulatta* lineage form *silenus/nigra* lineage (**supplementary Fig. S12**). The result upheld the topology displayed by the species tree (**Fig. 1B**). The phylogenetic tree based on introgressed and non-introgressed regions form *silenus/nigra* lineage resulted in different phylogenetic positions of the *fascicularis/mulatta* lineage.

### Population size history estimate

The history of the effective population size (*N_e_*) for macaque species were modeled from the distribution of heterozygous sites across the genome using a pairwise sequentially Markovian coalescent (PSMC) model [58] (**supplementary Fig. 13A-D** and **supplementary Fig. S14**). With the exception of *M. sylvanus* (**supplementary Fig. 13D**), which showed a truncated plot and most likely due to insufficient sequencing depth, the *N_e_*of remaining macaque species are very similar from 5 Mya until ~ 2.5 Mya, indicating that all Asian macaque species shared the same demographic history during the Pliocene. The Pliocene was a period of global cooling after the warmer Miocene and the cooling and drying of the global environment may have contributed to the enormous spread of savannas during this time [59]. The change in vegetation undoubtedly makes a huge impact on arboreal animals, such as macaques. Subsequently, the ancestor of most macaque species showed an upward trend of *N_e_* after ~2.5 Mya. However, the *N_e_* of three species of the *nigra* group remained stable until 1.5 Mya with a subsequent upward trend at different rates, possibly corresponding to population growth and expansion once ecological conditions allowed. Macaques except of the *nigra* group experienced a population growth until the Pleistocene glaciations and the *Ne* of different macaque species declined during different glaciations indicating that they likely were restricted to smaller refugia during glacial periods. During the last interglacial period, *N_e_* for *M. mulatta* (CR), *M. assamensis, M. nemestrina* and *M. tonkeana* increased rapidly but subsequently declined again.

## Discussion

### Phylogeny and classification of species groups

The present study enables us to verify the validity of species groups within the *Macaca* genus and to reconstruct their evolutionary history, which represents an exciting evolutionary scenery as a primate taxon whose “Out of Africa” spread mirrors that of non-African humans. The species tree deduced by ASTRAL supports the division of macaques into seven well-supported clades or lineages that correspond to the classification of the genus into seven species groups [21, 29] (**Fig. 1B**). *M. sylvanus* diverged first and *F_ST_* between *M. sylvanus* and the Asian macaques is higher than any pairs of Asian macaque groups (**supplementary table S6**). The results support the separation of *sylvanus* from the *silenus* group, which also confirmed the classification of Delson [23] based on morphology and geographic distribution. Asian macaques contained the *silenus* and *nigra* lineages and a clade including all other Asian lineages (**Fig. 1B**). The s*ilenus* and *nigra* lineages were grouped into a single species group (*silenus* group) by Fooden [22] and Delson [23], but recently separated into two distinct species groups: *silenus* and *nigra* or Sulawesi [21, 25, 29, 33]. Based on the species trees, the six investigated species of the *silenus* and *nigra* groups cluster into two clades: *silenus* groups (*M. leonina, M. silenus, M. nemestrina*) and *nigra* groups (*M. maura, M. tonkeana* and *M. nigra*) (**Fig. 1B**).

Except for the *silenus/nigra* lineage, the Asian macaques include two further clades. The first one includes *M. fascicularis, M. fuscata, M. cyclopis* and *M. mulatta* (**Fig. 1B**). Initially, all these four species were grouped into the *fascicularis* group [22], but Groves [25] separated *M. fuscata, M. cyclopis* and *M. mulatta* apart from *fascicularis* group into the *mulatta* group. The *F_ST_* values vary widely between *M. fascicularis* vs *M. mulatta* (0.171) and *M. fascicularis* vs *M. fuscata* (0.415). Yan, et al. [41] indicated that about 30% genome of mainland *M. fascicularis* are of Chinese *M. mulatta* origin, which shows that the lower genetic distance between *fascicularis* and *mulatta* groups is reasonable and likely caused by hybridization events. Therefore, they should still be divided into two different groups, particularly as *M. fascicularis* from the Sundaland region might not contain any genomic contribution of *M. mulatta*. The second clade contains *M. arctoides, M. assamensis, M. thibetana, M. leucogenys* and *M. sinica* (**Fig. 1B**). *M. arctoides* was assigned to the *arctoides* group [22], but based on molecular data, later integrated into the *sinica* group [23, 34, 35]. In contrast, Groves [25] recognized *M. arctoides* as member of the *fascicularis* group. Due to all these uncertainties, *M. arctoides* was recently separated into its own *arctoides* group [21, 29, 33].

### Contributions of ILS and ancient hybird to Gene Tree Heterogeneity

A recent study has reported gene flow events between the *fascicularis* and *silenus* groups and concluded that the introgression signals between these two groups are mainly due to genome similarity in closely related species [48]. In this study, we utilized QuIBL and discovered that among the 120 triplets examined, 60.8% exhibited phylogenetic discordances exclusively resulting from ILS. Moreover, evidence of introgression was only observed in 9.1% of the tested triplets (**Supplementary tables S8**), indicating that ILS rather than introgression is the primary factor contributing to the discordance among these species. Our research also suggests that ancient hybrid speciation may have occurred among the ancestral populations of *silenus/nigra* and *arctoides/sinica* clades. However, these analyses do not differentiate between ILS and introgression, both of which can create similar phylogenetic signals [60]. Thus, a plausible explanation is that *fascicularis/mulatta* clade may have originated from an ancient hybrid speciation of the two parental lineages (the ancestral populations *silenus/nigra* and *arctoides/sinica* clades) and is accompanied by ILS. The rationale is, first, according to PSMC analysis, all species in the genus *Macaca* were experiencing a population decline during the time window of the potential hybridization event (3.79 – 4.01 Mya). This decline of the ancestral population most likely resulted in the separation of subpopulations leading to rapid ILS [61]. Secondly, our *Dsuit* results based on two most common topologies (**Fig 3C** and **3D**), phylonet (**Supplementary tables S9-S10**) and HyDe results (**Fig. 3E** and **Supplementary tables S11**) all suggested a potential formation of a hybrid species. However, the authenticity of this event could be disturbed due to two reasons. First, it is difficult to detect the superposition of multiple introgression events throughout macaque history accurately. Additionally, ancient hybridization events may have unique evolutionary histories, such as ongoing gene flow, distinct introgression histories, maintenance of assortative mating, and rate heterogeneity [62]. Secondly, the power of HyDe and D-statistic in detecting hybridization events under high ILS and ancient rapid radiations is limited [63]. Simulation analyses have found that in the case of high level of ILS, the power of HyDe decreases to 0-0.1, and the power for the D-statistic was relatively poor either (0.03–0.7). Moreover, ancient hybridization can reduce the accuracy of the γ estimates [63]. However, though they are not the sole source, both ILS and potential hybrid speciation have contributed to the conflicts of macaques.

D-statistics also reveal asymmetric results for *fascicularis/mulatta* lineages which suggested introgression for *fascicularis/mulatta* lineages. In addition, PhyloNet and HyDe analyses suggested high levels of introgression or hybridization occurred for all macaques from *fascicularis/mulatta* lineage whether its current distribution is isolated or not. Furthermore, the phylogenetic tree based on SNVs from introgressed and non-introgressed regions, which was identified by *F*_d_ statistic and a two-layer hidden Markov model, resulted in different phylogenetic positions of the *fascicularis/mulatta* lineage. And the phylogenetic tree excluding SNVs located in all putatively introgressed regions for the *fascicularis/mulatta* lineage from *silenus/nigra* lineage (**supplementary Fig. S12**) upheld the topology displayed by the species tree (**Fig. 1B**). All these results suggested that potential ancient hybridization occurred in the *fascicularis/mulatta* lineage.

Ancient admixture of ancestral variation is a powerful means for rapid radiations to occur [64]. Hybridization analyses and QuIBL tests support this idea. However, apart from ancient introgression between macaque groups, secondary introgression and recent gene flow can also result in differences in D-statistics. Though HyDe and the D-statistics are powerful for detecting hybridization in most of scenarios [63], to what extent the ILS or potential hybrid have shaped the macaque history may require large-scale genomic data sets to perform optimally under complex evolutionary scenarios in future.

### The radiation of macaques

Fossil data suggest that the genus *Macaca* arose about 7 Mya in North Africa during the Late Miocene [23, 59, 60, 61], which is consistent with our MCMCTree result (7.76 Mya, **Fig. 2**). The Messinian Salinity Crisis (5.9-5.3 Mya), in which the level of the Mediterranean Sea fluctuated and occasionally dried out, may have facilitated macaque dispersal from North Africa [51]. A large number of macaque fossils have been reported in Europe near the Mediterranean [33, 51]. Macaque ancestors likely invaded Europe via the arid Mediterranean during Messinian stage [26, 33, 46, 51]. The ancestor of *M. sylvanus* split from the main stem ~5.29 Mya and settled around the Mediterranean (**Fig. 2** and **Fig. 4**). Dental specimens of the oldest known Asian fossil referred to the genus *Macaca* are *M. palaeindicus* from the Late Pliocene Tatrot formation of the Siwalik Hills [23, 59, 62]. Hence, the ancestor of Asian macaques spread eastward step by step likely along the Siwalik Hills to the Hengduan Mountains, which are a potential diversification hotspot for primates [1, 65, 66, 67, 68], and diverged into different lineages. Firstly, the ancestor of *silenus/nigra* split ~4.01 Mya and then hybridized with the ancestor of other Asian macaques and formed a pattern of three clades (**Fig. 4**).

**Figure 4.**
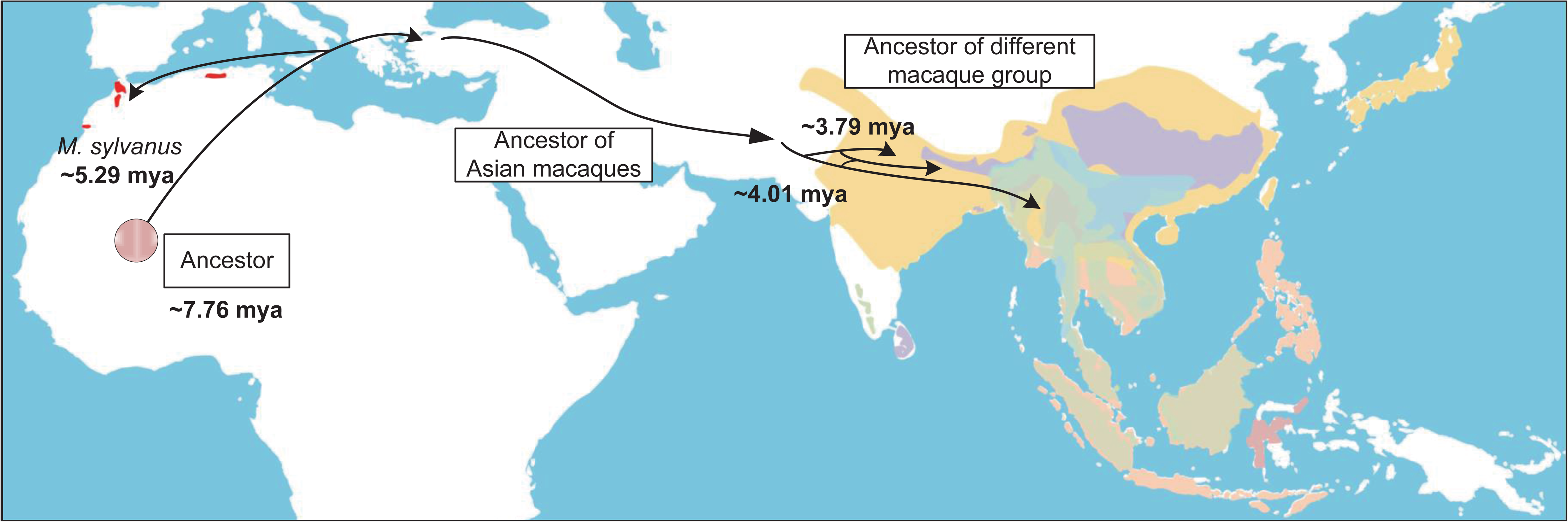
Scenario for the radiation of macaques. Macaques originated in Africa and after the split of *M. sylvanus* the ancestor of Asian macaques invaded Asia and initially split into three lineages, *silenus/nigra*, *arctoides/sinica* and *fascicularis/mulatta*. *fascicularis/mulatta* clade may have originated from an ancient hybrid speciation of the two parental lineages (the ancestral populations *silenus/nigra* and *arctoides/sinica* clades) at about 3.79 – 4.01 Mya ago.

## Conclusions

In this study, we conducted a phylogenetic analysis of 24 whole macaque genomes of which 16 are newly generated. We identified ~20 million high-quality SNVs in each macaque individual, which can be invoked as important baseline data in other studies. We support the division of macaques into seven species groups and found that there were visible differences in *F_ST_* within and between species groups. Importantly, our results reveal *fascicularis/mulatta* clade may have originated from an ancient hybrid speciation of the two parental lineages (the ancestral populations *silenus/nigra* and *arctoides/sinica* clades) and is accompanied by high levels of ILS. Most prominently, we have used the available fossil data and the literature to reconstruct the radiation scenario of the genus macaques. In summary, this study suggests that high level of ILS and potential hybrid speciation play important roles in the radiation of macaques.

## Materials and methods

### Sample collection and library preparation

Samples of 16 macaque individuals were obtained from the Institute of Zoology, Chinese Academic of Sciences and the German Primate Center (**supplementary table S1**). In addition, genome data of eight other macaque individuals and one *Papio anubis* individual were obtained from the NCBI Short Read Archive (accession no. show in **supplementary table S1**). The present study was approved by the Animal Ethics Committee of the Institute of Zoology, Chinese Academy of Sciences. The procedure of blood and tissue collection was in strict accordance with the Animal Ethics Procedures and Guidelines of the People’s Republic of China. DNA was isolated from blood or tissue samples with the Qiagen Blood and Tissue Kit. Sequencing libraries were prepared based on KAPA library preparation kit with insert sizes of 300-500 bp and sequenced using Illumina Hi-Seq 2500 and 4000 technologies. Quality control was performed using FastQC with default parameters (www.bioinformatics.babraham.ac.uk/projects/fastqc/).

### Data Processing

Paired-end reads for all samples were mapped to the *Papio anubis* reference genome (Panu_3.0, GCA_000264685.2) with Burrows-Wheeler Aligner (BWA) mem v0.7.17 [69], except for *M. nemestrina* (2), which was trimmed by cutadapt.2.4 [70] firstly because of the poor sequencing quality due to DNA degradation. The output bam files were sorted using samtools [71] and duplicates were marked with GATK-v4.1.2.0 [72]. Single-nucleotide variants (SNVs) were called using GATK 4.1 [72] following best practice. We obtained the GVCF file for each individual using the “HaplotypeCaller” method in GATK and then all samples were merged based on “CombineGVCFs”. We used the GenotypeGVCFs-based method with the “includeNonVariantSites” flag to get the vcf files. After that, we applied the “SelectVariants” to exclude indels and split the variant and nonvariant sites. All nonvariant sites were filtered in subsequent analyses. We applied the hard filter command “VariantFiltration” to exclude potential false-positive variant calls with the following criteria: “-filterExpression QD< 5.0 || FS > 60.0 || MQ < 40.0 || ReadPosRankSum< −8.0 || MQRankSum < −12.5” and “-genotypeFilterExpression DP < 4.0” [73]. All variant calls that failed the filters were removed. All SNVs were annotated by ANNOVAR v2013–06-21 [74]. (**supplementary tables S3** and **S4**) based on gff3 file from Ensembl (Papio_anubis.Panu_3.0.104.gff3). Heterozygosity is defined as the rate of heterozygous SNVs in the genome, which were further filter by the following parameters: “--minGQ 20” and “--minDP 8” to ensure data quality based on VCFtools. We calculated the heterozygosity rate in a nonoverlapping window (window = 100 kb) for all 24 macaques. Genome-wide distribution of heterozygosity was shown in a box plot using boxplot in R package (**supplementary table S2** and **supplementary Fig. S2**).

### Phylogenetic reconstruction using the window-based strategy

We constructed the consensus sequences for all individual genomes and aligned them per chromosome (sex chromosomes, mitogenomes and scaffolds were excluded) based on VCF, which did not include variants present in the Papio reference but were invariant among macaques. In addition, indels, repetitive regions and low-quality regions were removed from the consensus sequences. Per-chromosome alignments were split into nonoverlapping windows of 200 kbp. The maximum likelihood (ML) trees for each window were performed with RAxML v8 [75] with 100 bootstrap replicates and the GTR + CAT substitution model. From all 200 kbp window trees, we reconstructed a species tree with ASTRAL v4.10.5 [76] under the multispecies coalescent model returning quartet scores and posterior probabilities. The species tree was rooted with *Papio anubis*. Consensus networks of the window trees were generated using SplitsTree5 [40] with different thresholds. Genealogical discordance across the genome of rhesus macaque was assessed by Twisst [77]. All GF trees were clustered based on PhyBin (https://github.com/rrnewton/PhyBin) and the top two topologies were visualized by DensiTree [78].

### Phylogenetic analysis of concatenated whole-genome SNV data

SNVs were extracted from the filtered VCF file and converted into individual FASTA sequences using a python script (https://doi.org/10.5281/zenodo.2540861). Merging all FASTA sequences into a single file provided a multiple sequence alignment of all individuals and all concatenated SNVs. Phylogenetic trees were reconstructed with maximum-likelihood (ML) using RAxML v8 [75] with 100 bootstrap replicates and the GTR + CAT substitution model. In addition, a neighbor-joining (NJ) tree was constructed using the program TreeBeST V.1.9.2 (http://treesoft.sourceforge.net/treebest.shtml), which has a build-in algorithm to build the best tree that reconciled with the species tree and rooted with the minimized number of duplications and losses. To further visualize genetic relationships among macaques, we performed principal components analysis (PCA) on the filtered autosomal SNVs using the Eigensoft package (version 5.0) [79].

### Phylogeny of whole mitochondrial genomes

We used NOVOPlasty [80] to *de novo* reconstruct the mitochondrial genomes (mitogenomes) for 18 macaques representing 15 species (**supplementary table S5**) and then annotated the genome sequences using MitoZ [81]. In addition, we downloaded 15 mitogenomes for the genus *Macaca* (**supplementary table S5**) from NCBI based on the filter criteria mentioned in Roos, et al. [33]. The 33 mitogenomes were aligned using MAFFT version 7 [82] and indels and poorly aligned positions were removed with Gblocks version 0.91b [83]. A ML tree was reconstructed using RAxML v 8 [75] with a GTR + CAT substitution model and applying 1000 bootstrap replicates to verify nodal support.

### QuIBL

We used QuIBL [18] to assess the likelihood of introgression and ILS for each locus in every species triplet under evaluation. QuIBL provides estimates of the proportion of introgression and the probability of a locus falling into a model with either introgression or ILS alone. To begin, QuIBL estimated the internal branch length distribution for each locus in the triplet of species. Given its sensitivity to recombination [18], we extracted 20-kb windows separated by 400-kb from each sample to minimize the risk of including a recombination breakpoint in the window. We filtered the inferred ML trees based on the number of parsimony-informative sites (≧10), resulting in a total of 3,175 trees that served as QuIBL inputs. We used the species tree topology to assign the outgroup to each triplet and also calculated the percentage of loci that supported discordant topologies and showed significant evidence for introgression.

In order to determine whether ILS (scenario 1) or a combination of ILS and introgression reflected the inner branch lengths, QuIBL evaluated the probability values (Bayesian Information Criterion test, BIC) (scenario 2). The scenario of ILS only with the lower BIC value is preferred when Delta. BIC (BIC_scenario2_-BIC_scenario1_) is larger than 10 and the BIC value is less than 0. The combination of ILS and introgression with the lower BIC value, however, is preferred when Delta.BIC is smaller than −10.

### D-statistics

We implemented D-statistics with *Dsuite* [53] across all combinations of the 16 macaques described above. The topology required for *Dsuite* was consistent with two alternative topologies described in **Fig. 3A-B**, respectively. Then, the D-statistics of all possible combinations with *Papio anubis* as the outgroup (Out) and three different macaque species with a phylogenetic topology following (((M1, M2), M3), Out) were calculated with the Dtrios module. In total, 566 groupings were analyzed for each topology. After that, D-statistics were calculated for each branch of the alternative topologies using the f-branch module, and visualized the statistical results using dtools.py script provided in the *Dsuite* software.

### Hybridization Detection

To obtain further support for the ancient introgression suggested by the above approach based on the D-statistics, we applied HyDe (Hybridization Detection) [55], which can automate the detection of hybridization across large numbers of species despite ILS and can perform hypothesis tests at the population or individual level by estimating the amount of admixture (γ). The detailed analysis procedure is as follows: We applied the population approach in “run_hyde_mp.py” script to test hybridization events on all triples among macaque lineages in **Fig. 1C** (*fascicularis*/*mulatta* clade, *arctoides/sinica* clade, *silenus/nigra* clade, and *sylvanus* clade) with *Papio anubis* as the outgroup. Results are first filtered based on whether there are significant *P* values (*P* values <0.05) for hybridization. In addition, different γ values represent different events. For example, a 50:50 hybrid is characterized by a γ value of about 0.5 and a very low levels of admixture (e.g., 0.01 means close to parent P1 while 0.99 means close to parent P2) can be indicators of different processes such as ILS and more ancient hybridization. We found that a 50:50 hybrid was significantly concentrated among the *silenus/nigra*, *fascicularis*/*mulatta*, and *arctoides/sinica* clades. We then used the script “run_hyde_mp.py” to test each individual within a putative hybrid population (*fascicularis*/*mulatta* clade) using specified triples and then performed bootstrap resampling (500 replicates) of the individuals within the putative hybrid lineages for each specified triple based on “bootstrap_hyde.py” script.

### Maximum likelihood inference for reticulation with PhyloNet

We used the InferNetwork_MPL program in PhyloNet [54], which is based on Maximum Pseudo Likelihood (MPL) algorithm, to analyze a set of every 3th GF ML tree, that is, 2150 trees in a coalescent framework that accounts for incomplete lineage sorting (ILS) while allowing different numbers of reticulation events. In addition, the individual “2” in all macaque species were pruned from the input gene trees to reduce complexity and computational demand. We set the numbers of reticulation events as 1 and 2 to conduct two independent analyses and each analysis was run with 100 iterations, yielding the five networks with the highest likelihood scores. We visualized the optimal networks in Dendroscope v3 [84] and present it in **supplementary Fig. S10.**

### *F*_d_ value

To identify putatively introgressed regions in macaques, we screened these segments according to the strategy of Martin, et al. [56] and van der Valk, et al. [6]. The method predicts that introgression between species at a specific genome region should reduce the between-species divergence in this region as compared to the rest of the genome [6, 56]. We computed the *F*_d_ value, which is sensitive to the number of informative sites within windows, for each 10-kb window across the whole genome of each macaque.

### Introgression analysis with efficient local ancestry inference

The two-layer admixture model implemented in efficient local ancestry inference [57] was used to infer introgressed segments in *fascicularis/mulatta* (the admixture population, -p 1) with the *silenus/nigra* as source population 2 (-p 11) and *arctoides*/*sinica* were used as source population 1 (-p 10). We run 30 iterations with mixture generation values of 2,910,000, 3,200,000 and 3,450,000, which was estimated based on MCMCTree result, and the averaged results over the three independent runs were used for detecting introgression tracts. Sites with a proportion of source population 2 ancestry greater than 1.5 were defined as introgression sites from *silenus/nigra* lineage which may cause position conflict of *fascicularis/mulatta* lineage in species tree. We sum the proportion of source population 2 ancestry in each 10 kb windows, which is same to *F_d_* and the top 1% windows were defined as putatively introgression tracts.

### Analysis of demographic history using genome-wide data

We applied the pairwise sequentially Markovian coalescent (PSMC, v0.6.5-r67) analysis [58] to estimate the demographic history of each macaque individual (**supplementary Fig. S2**). We use the Samtools mpileup [71] to generate the input files for PSMC with the parameter “-C” set to be 50 to improve mapping quality. In addition, only autosomal SNVs were reserved to generate the diploid sequence for PSMC. PSMC was run with 25 iterations (-N), a maximum 2N0 coalescent time (-t) of 15, an initial theta/rho ratio (-r) of 5, and the 64 time-intervals were parameterized as “4 + 25*2 + 4 + 6”. PSMC plots were scaled with a mutation rate (μ) of 0.9×10^-8^ and a generation time (g) of 11 years for all macaque species [85]. Bootstrap analyses (100 bootstrap replications) were performed for each species (**supplementary Fig. S14**).

### Divergence time calibration

Coding sequences (CDSs) and protein sequences for different representative outgroup species and three macaques (*M. mulatta, M. fascicularis, M. nemestrina, Papio anubis, Homo sapiens, Pan troglodytes* and *Pongo abelii*) were retrieved from Ensembl (release-98). The single-copy orthologue gene list shared by these species was obtained using OrthoFinder [86]. The macaques were mapped to the Panu_3.0 genome and thus their CDSs have the same genomic coordinates. Therefore, the CDSs of all macaques were extracted after ortholog detection. CDSs of these single-copy genes were aligned by PRANK (v.170427) [87] under codon model and concatenated into a single super-matrix. Fourfold degenerate (4D) sites were extracted from them using MEGA7 [88] and concatenated to calibrated the divergence time of the genus *Macaca* with MCMCTree program in PAML v4.9 (Phylogenetic Analysis of ML, v4.9) package [49]. Four calibration points were applied to constrain the divergence time of the nodes: Catarrhini 20–38 Ma, Hominidae 13–18 Ma, *Homo-Pan* 6–7 Ma and *Papio*-*Macaca* 5–8 Ma [33, 89]. To check for convergence of the stationary distribution, the analysis was run in duplicate, and the results were compared between runs and similar results for divergence time of macaques were obtained (**supplementary Fig. S9**).

## Acknowledgements

The authers thank Prof. Luo Shujin and Dr. Sun Xin for their assistance in DNA isolation, library preparation and quality control for *M. nemestrina (2)*.

## Author contributions

ML conceived this study, and X.Z. and C.R. contributed to the idea for this study. X.T., J.Q. and Z.L. performed experiments, analyzed data and wrote the draft manuscript. G.L., L.Z. and Y. S. help to analyze data. P.F. C.R. and J.L. provided samples. M.L., X.M. and C.R. revised the draft manuscript. All authors contributed to the final text.

## Data accessibility

Whole genome resequencing data produced in this study were deposited at the NCBI Sequence Read Archive (SRA) under Bioproject PRJNA732172.

