## Supplementary figures for "Phylogenomics reveals incomplete lineage sorting and ancient hybrid drove the radiation of macaques"

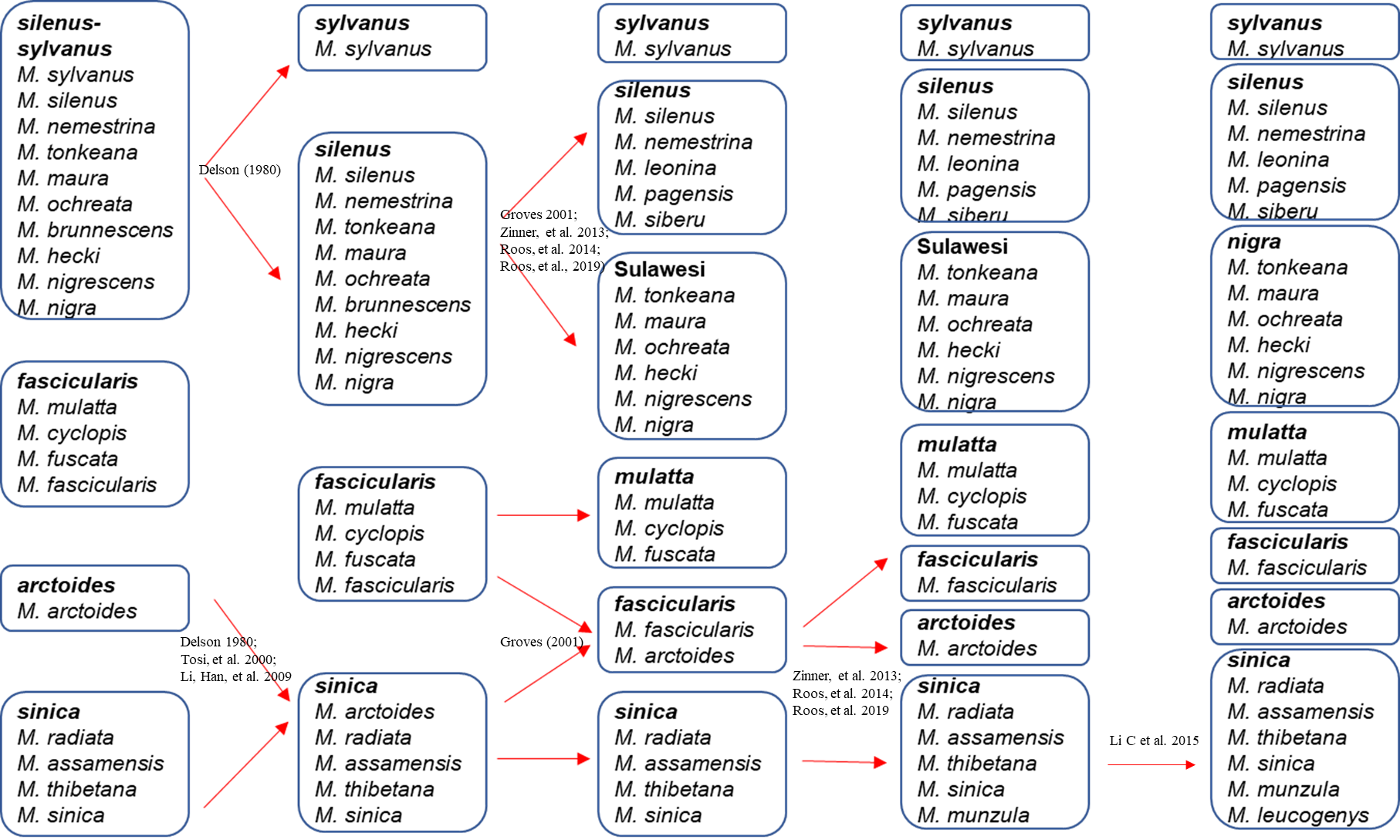
Figure S1. Changes in the classification of extant macaque species after Fooden (1976).

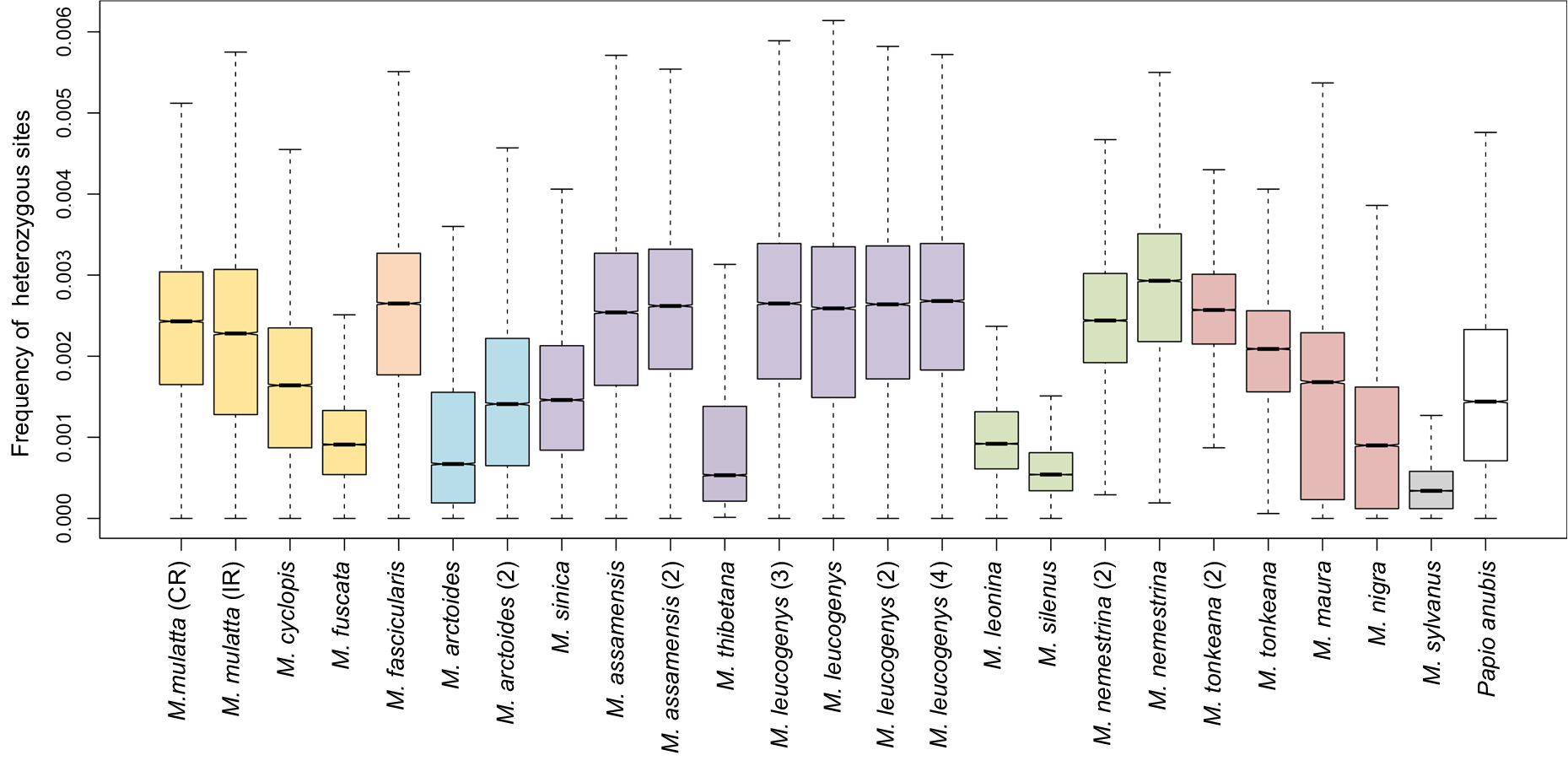

Figure S2. Genome-wide heterozygosity estimated from genomic 100-kbp windows.

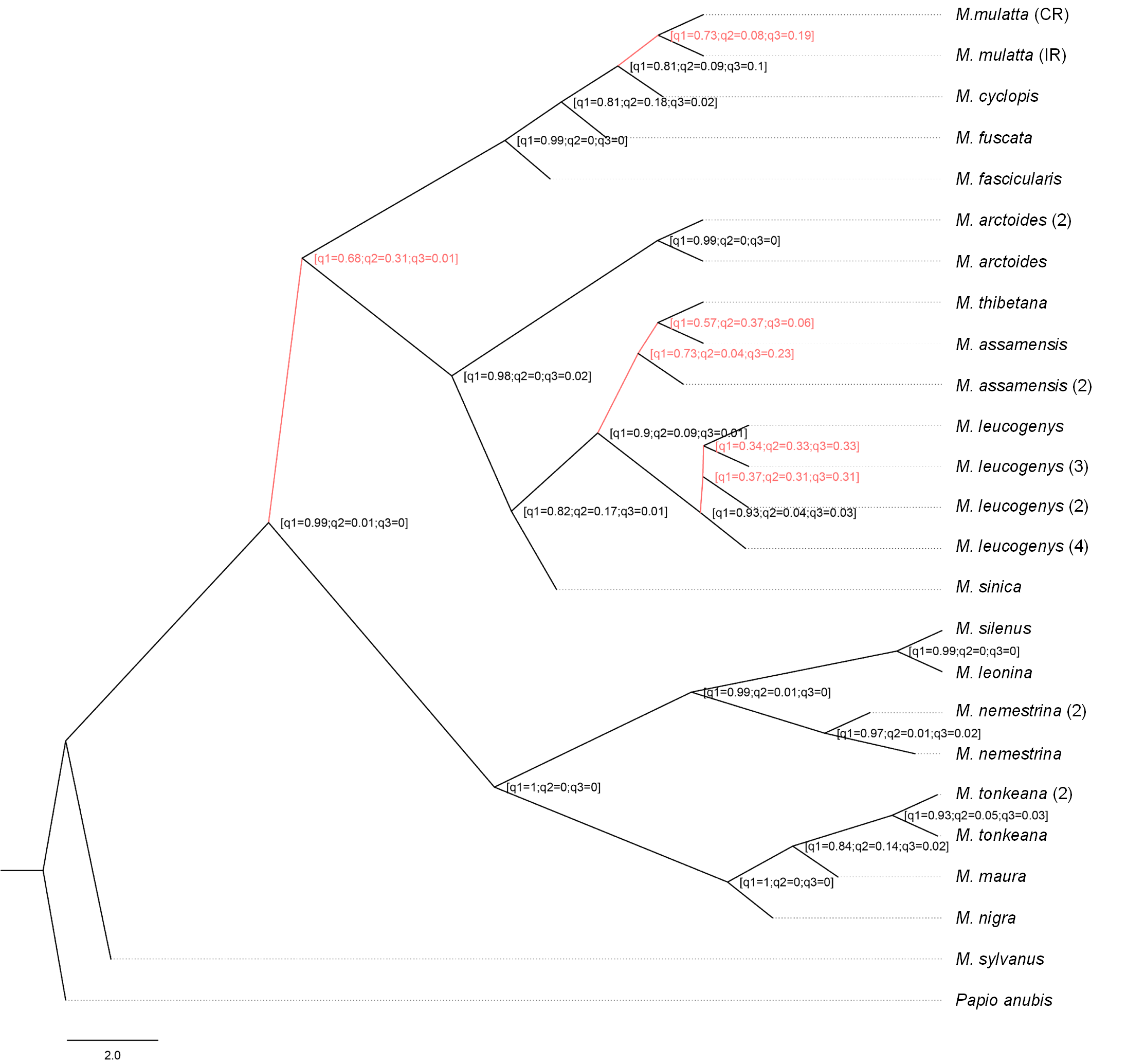

Figure S3. MSC-based species trees generated by ASTRAL using genome fragments. The tree was rooted with *Papio anubis*. ASTRAL quartet-scores for all branches are shown and branches labelled in Figure 1B are showed in red.

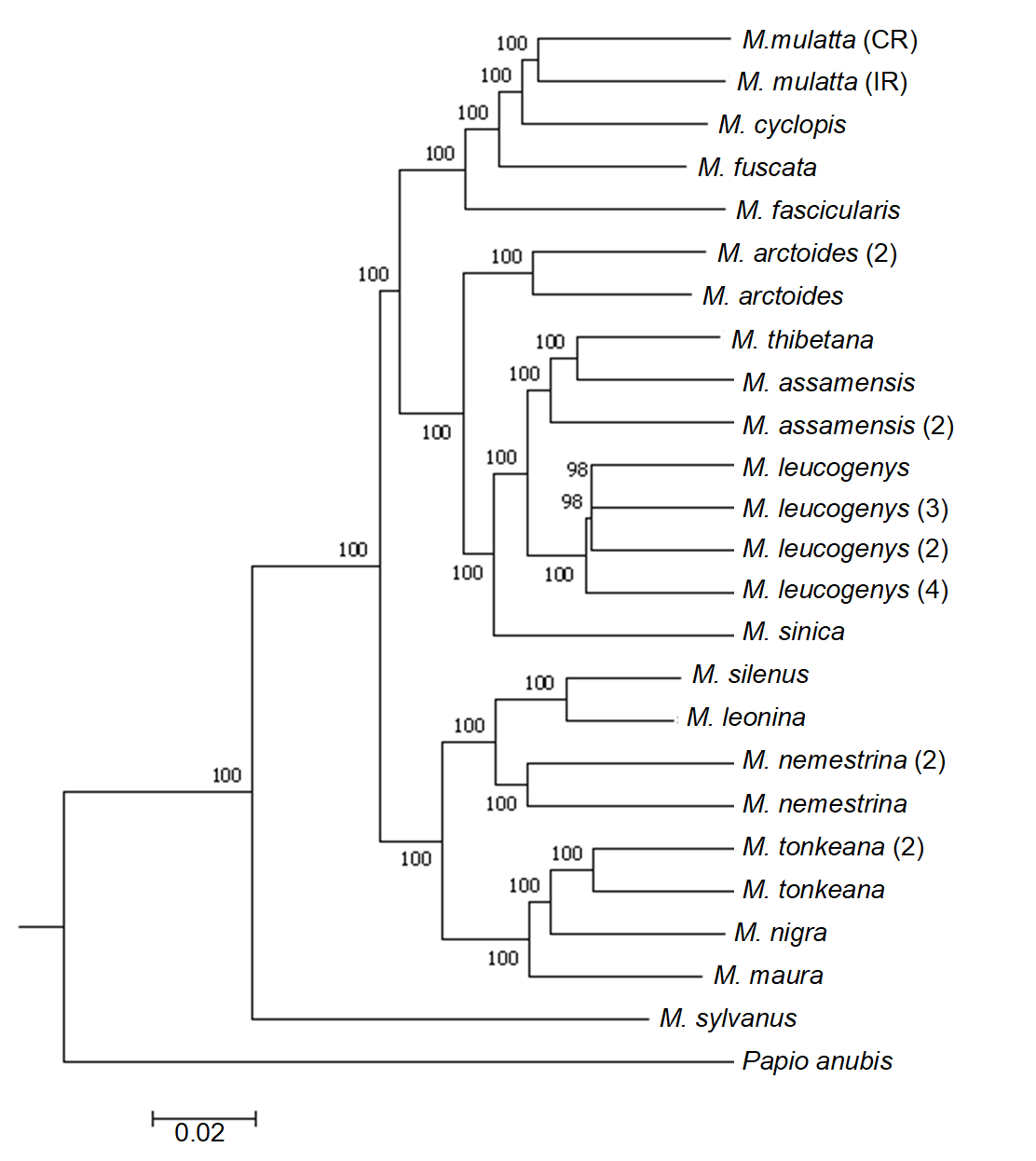

Figure S4. NJ tree based on Treebest using autosomal SNVs. Bootstrap values are given at nodes.

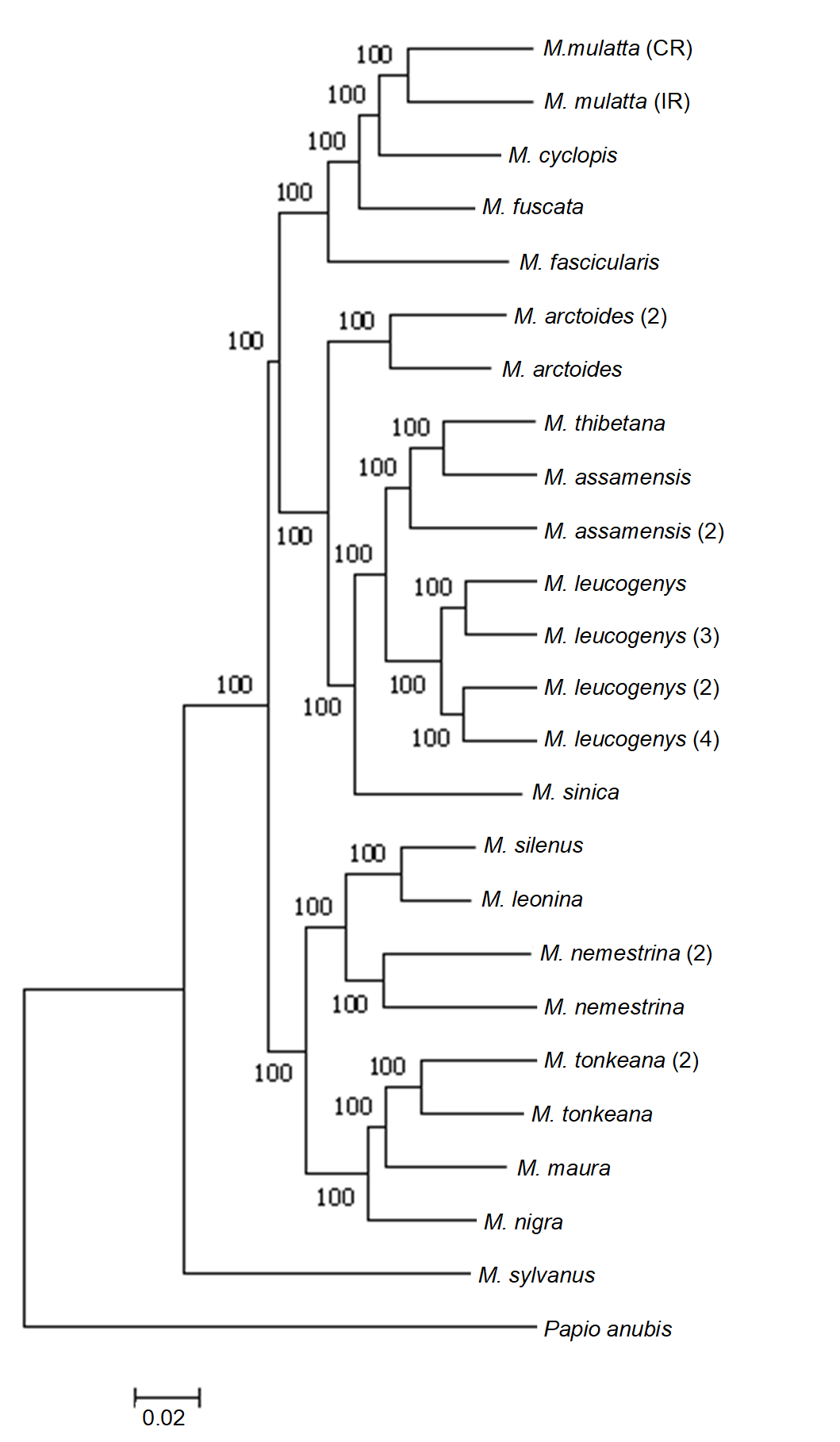

Figure S5. ML tree based on raxml using autosomal SNVs. Bootstrap values are given at nodes.

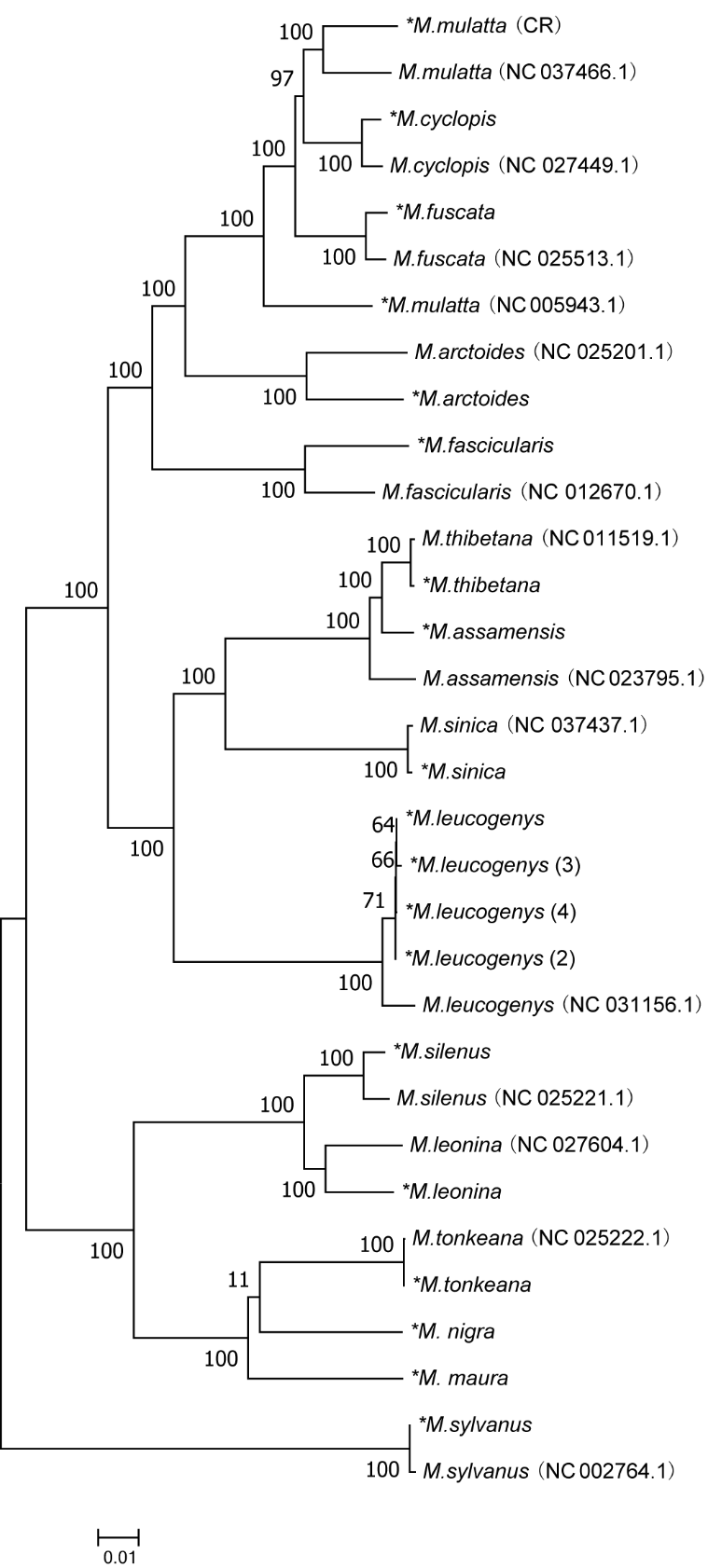

Figure S6. ML tree based on RAxML using mitogenomes. Newly generated sequences are marked with an asterisk. Accession numbers of published sequences are given in parentheses. Bootstrap values are given at nodes.

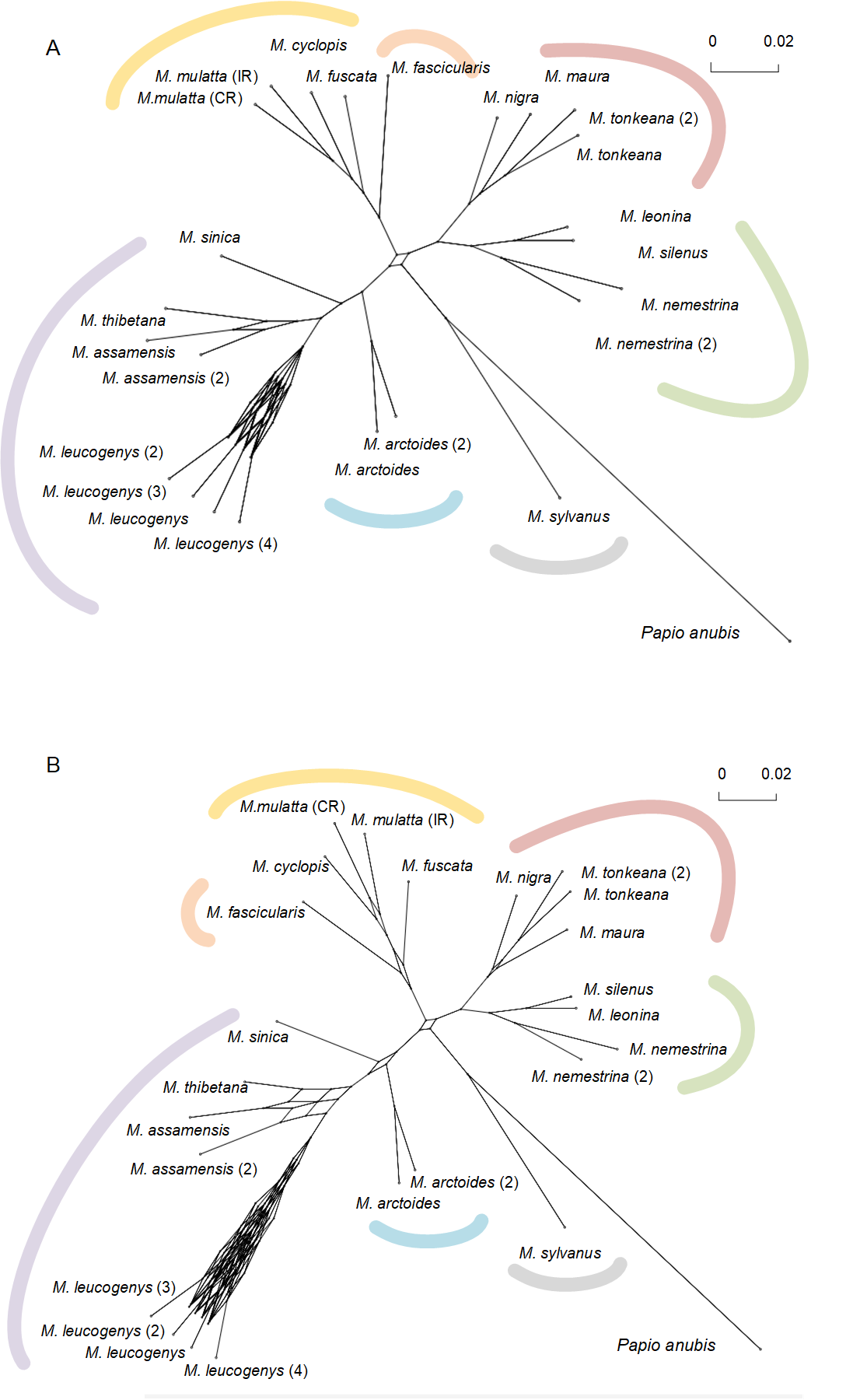

Figure S7. Consensus networks for macaques from 6,351 gene trees (200-kbp genome fragment) at different minimum thresholds (A: minimum threshold=20%; B: minimum threshold=10%) of gene trees to form an edge.

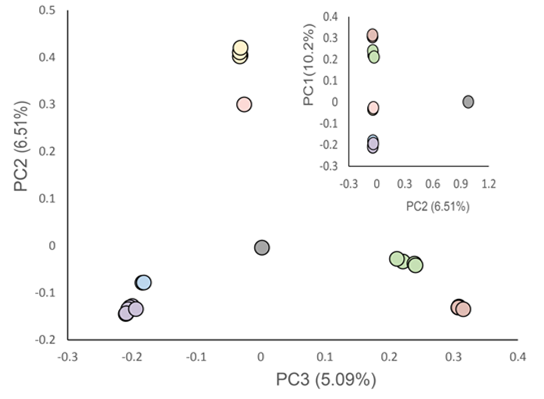

Figure S8. Plots of principal components 1 - 3 from PCA analysis of 24 macaques using autosomal SNVs.

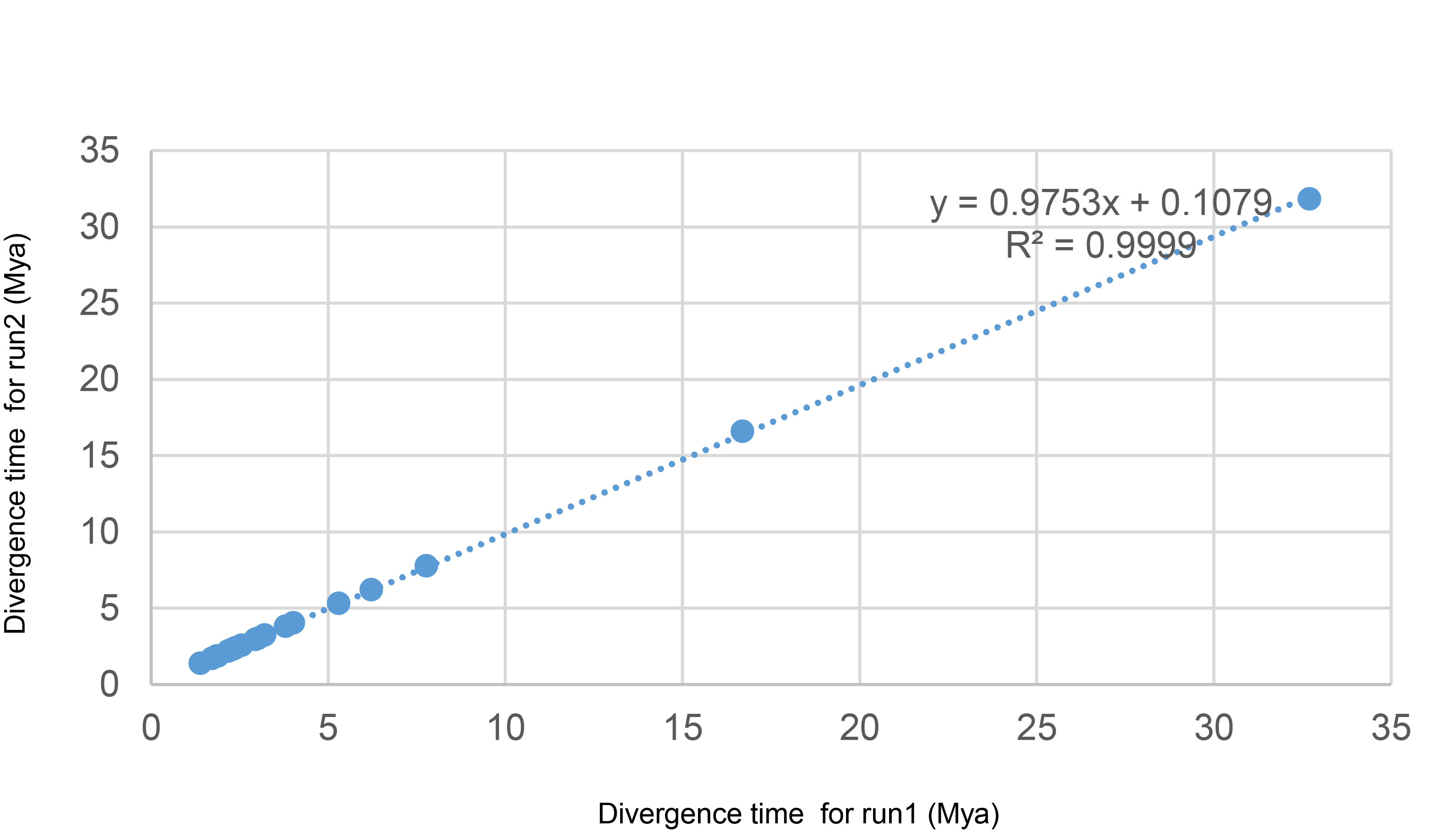

Figure S9. Convergence of the distribution for MCMCtree results.

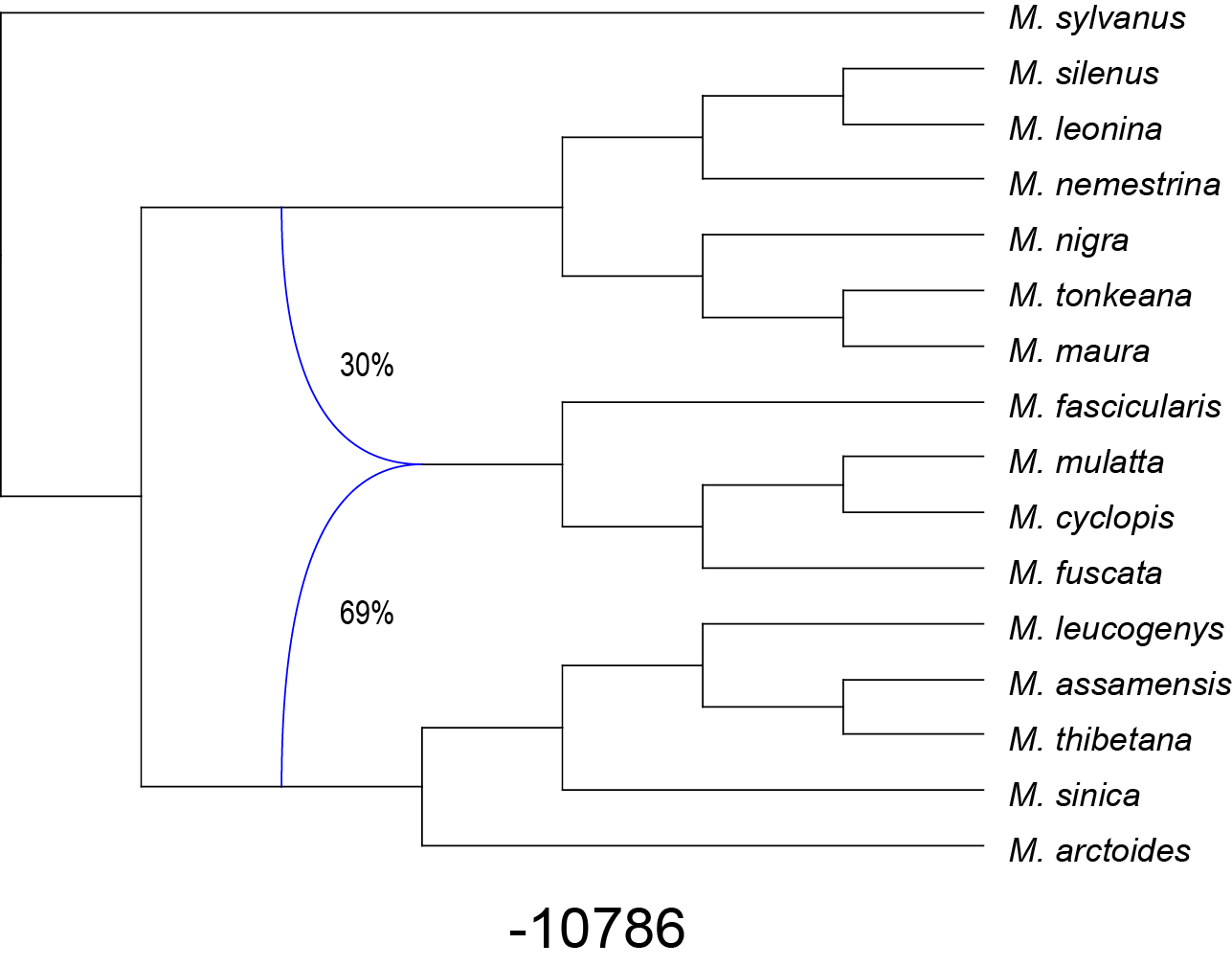

Figure S10. Rooted networks for the *Macaca* phylogeny with one reticulation inferred from PhyloNet. Reticulations are shown as blue arrows with inheritance probability denoted. Log-likelihood scores are shown below the networks. Notably, inheritance probability around 30% between *silenus*/*nigra* lineage and *fascicularis*/*mulatta* lineage resembles the distribution of quartet scores and the phylogenetic signals from genome fragments (Fig. 1).

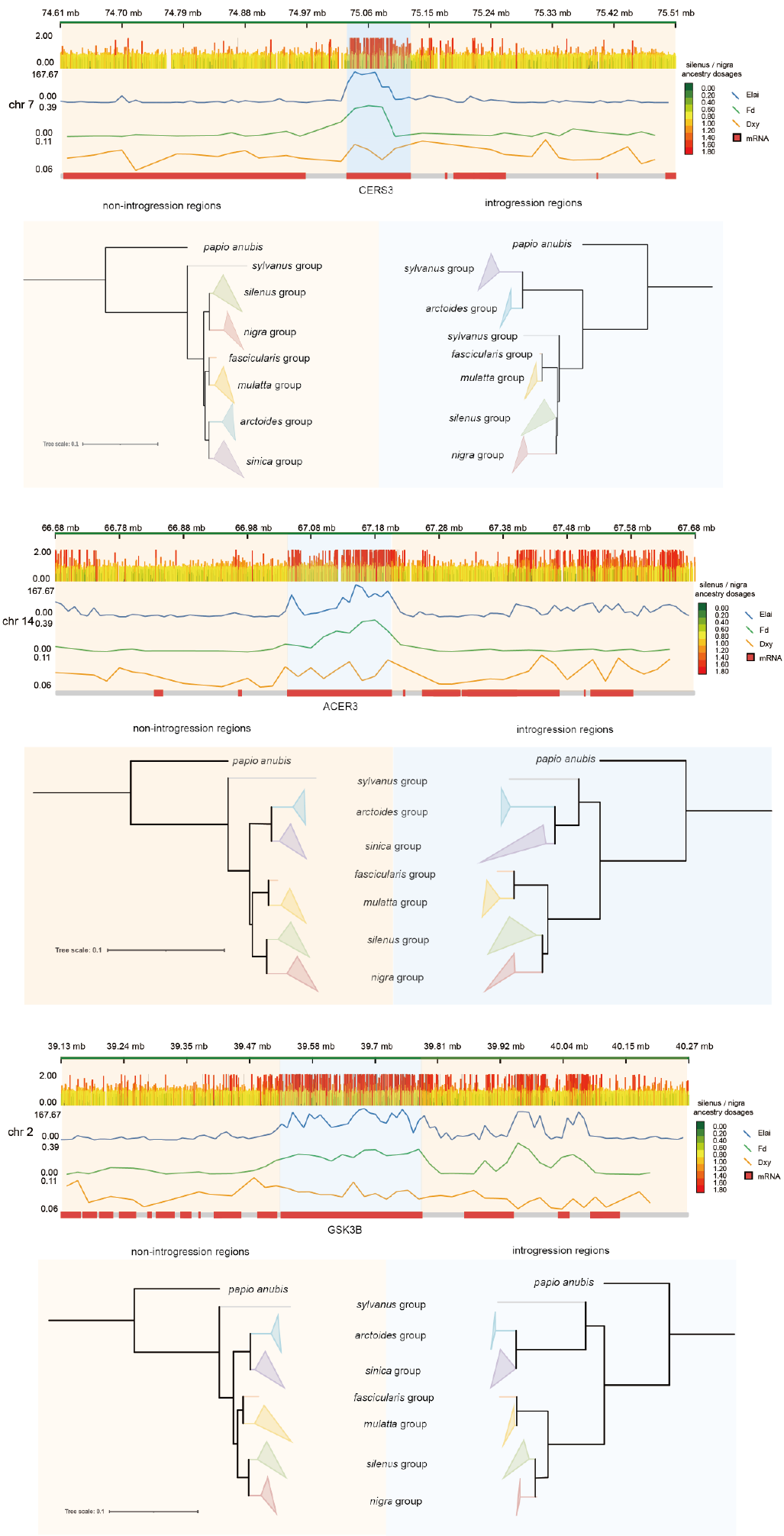

Figure S11. The schematic for inferred of local ancestry for *fascicularis/mulatta* lineage. First line showed 1 Mb regions which contains introgressed windows. The second part, which is a heatmap show the ancestry dosages for *fascicularis/mulatta* source from *silenus/nigra* lineage. Sites with a proportion of source population (*silenus/nigra* lineage) ancestry greater than 1.5 were defined as introgression sites. The third line show the average value of a proportion of source population (*silenus/nigra* lineage) ancestry in 10 kb windows. The fourth and fifth lines show the *Fd* and *Dxy* and the last part showed the genes in this region. The phylogenetic trees at the bottom of the figure showed the topologies based on the introgression windows and the non-introgression regions within this 1 Mb regions.

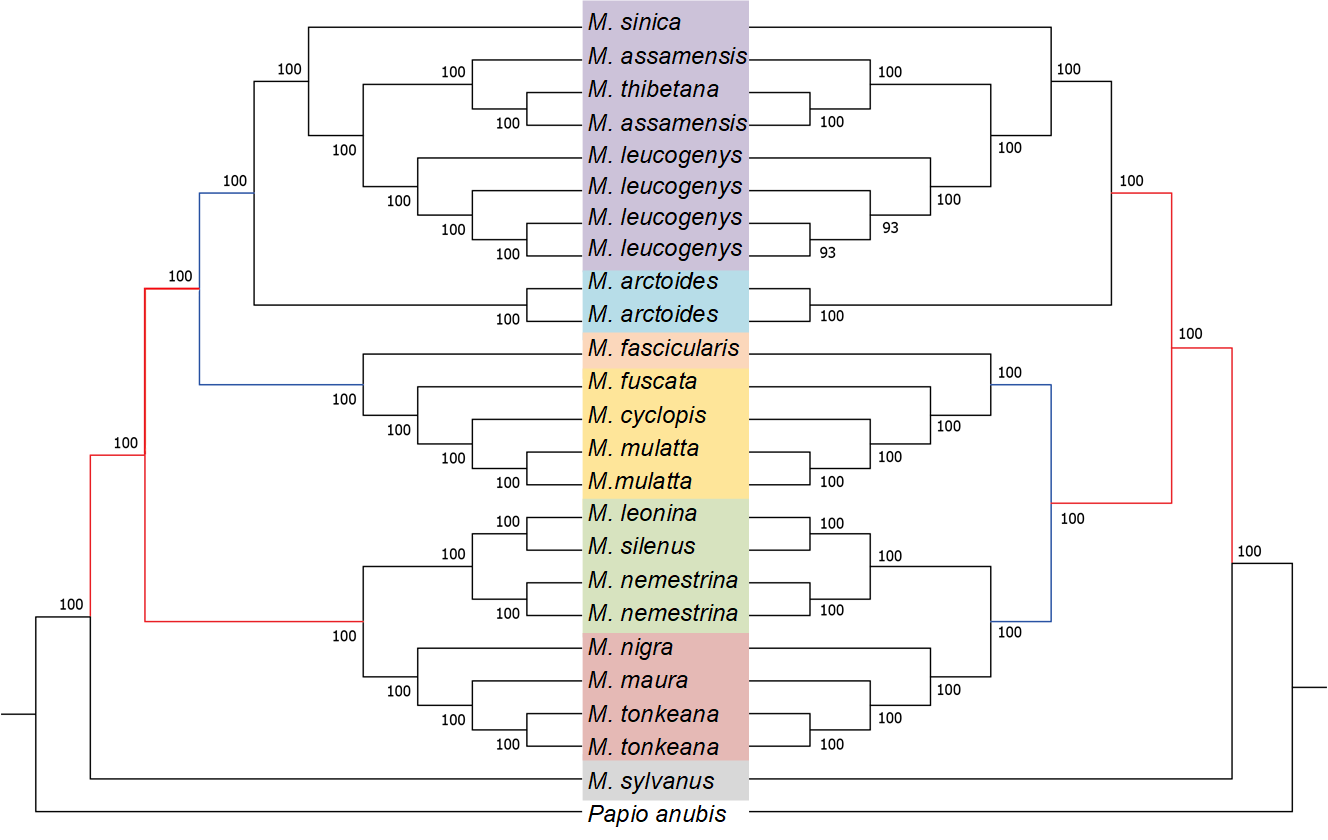

Figure S12. ML tree based autosomal SNVs located in introgressed windows and ML tree based on autosomal SNVs except those SNVs located in introgressed windows between the ancestors of *fascicularis/mulatta* and *nigra/silenus*. Bootstrap values are given at nodes.

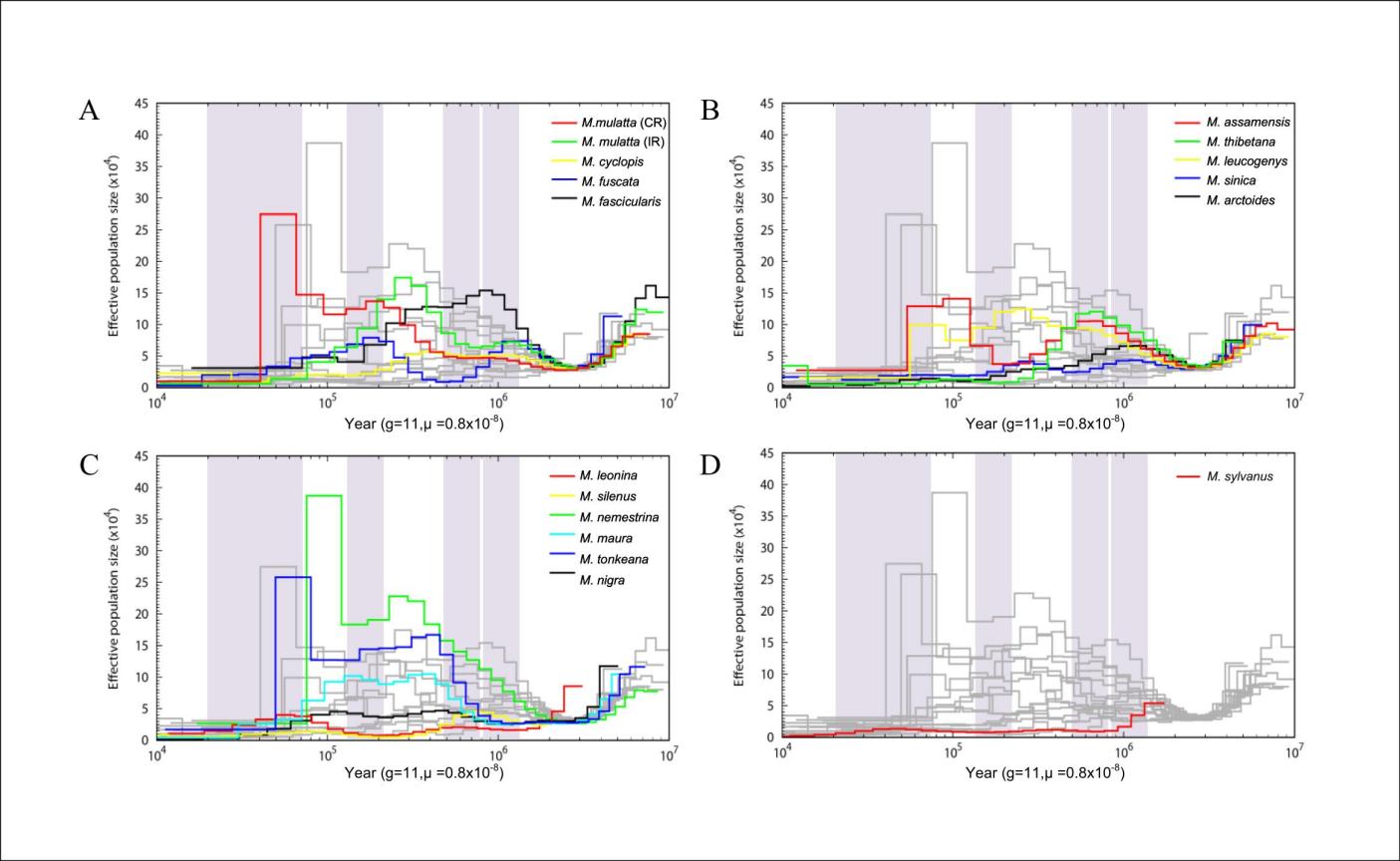

Figure S13 Historical effective population size.

(A-D) Historical effective population size using PSMC analyses for all macaque genomes. The x axis shows the time, and the y axis shows effective population size. Plots were scaled using a mutation rate (m) of 0.8 × 10^− 8^ substitutions per nucleotide per generation and a generation time (g) of 11 years. Light purple shading indicates glaciation in the Pleistocene and Holocene: Xixiabangma Glaciation (XG, 1.17~0.8 Mya), Naynayxungla Glaciation (NG, 0.78~0.50 Mya), Penultimate Glaciation (PG, 200–130 kya) and Last Glaciation (LG; 70–10 kya)

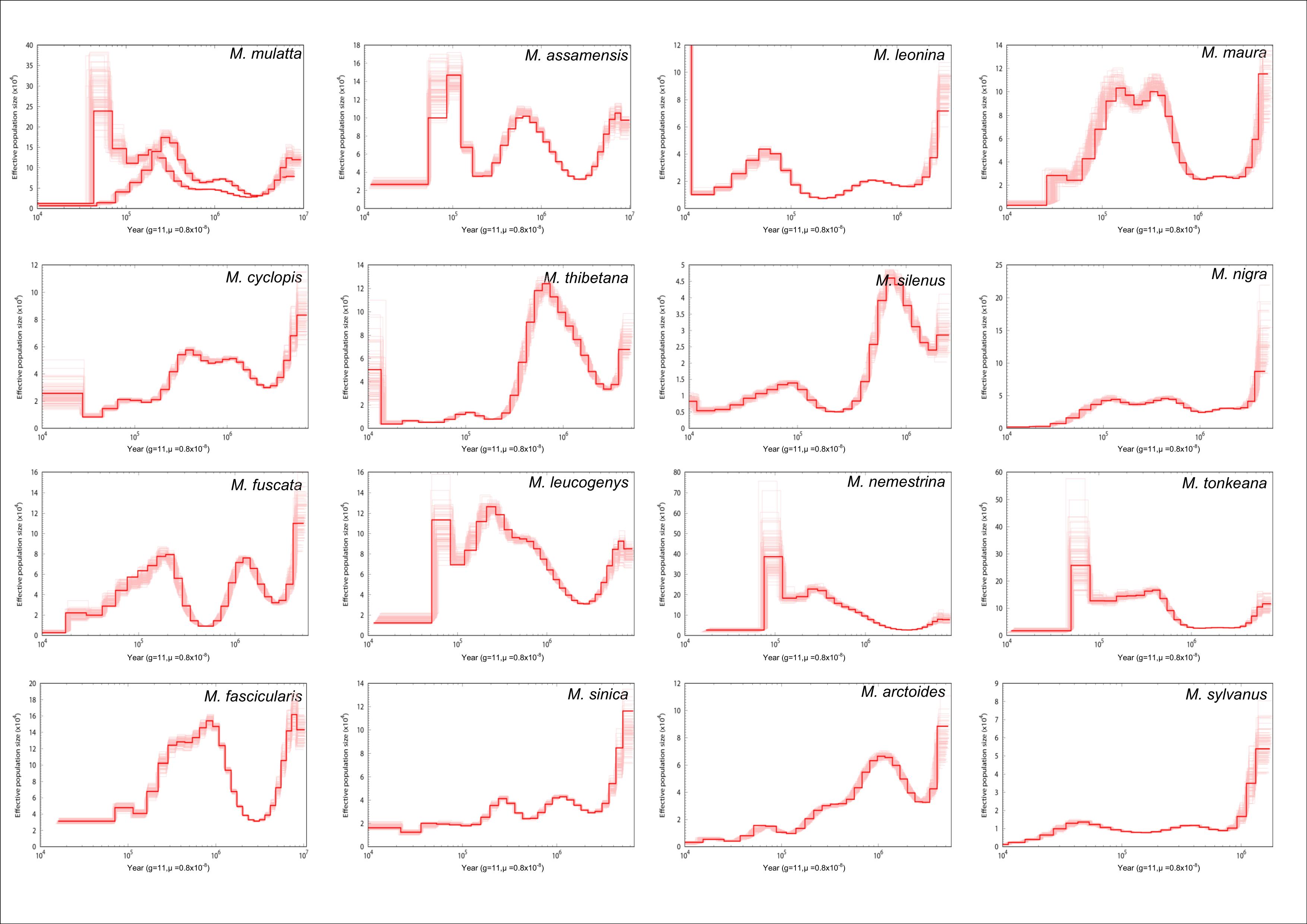

Figure S14. Demographic histories for each individual macaque genome with 100 bootstrap replicates. Each panel shows estimated ancestral effective population size calculated by PSMC. Bootstrap replicates are shown in light red. Sample names are given above each panel.
